# The preservative sorbic acid targets respiration, explaining the resistance of fermentative spoilage-yeast species

**DOI:** 10.1101/2020.04.09.034074

**Authors:** M. Stratford, C. Vallières, I.A. Geoghegan, D.B. Archer, S.V. Avery

## Abstract

A small number (10-20) of yeast species cause major spoilage in foods. Spoilage yeasts of soft drinks are resistant to preservatives like sorbic acid and they are highly fermentative, generating large amounts of carbon dioxide gas. Conversely, many yeast species derive energy from respiration only and most of these are sorbic acid-sensitive, so prevented from causing spoilage. This led us to hypothesize that sorbic acid may specifically inhibit respiration. Tests with respiro-fermentative yeasts showed that sorbic acid was more inhibitory to both *Saccharomyces cerevisiae* and *Zygosaccharomyces bailii* during respiration (of glycerol) compared with fermentation (of glucose). The respiration-only species *Rhodotorula glutinis* was equally sensitive when growing on either carbon source, suggesting that ability to ferment glucose specifically enables sorbic acid-resistant growth. Sorbic acid inhibited the respiration process more strongly than fermentation. We present a dataset supporting a correlation between the level of fermentation and sorbic acid resistance across 191 yeast species. Other weak acids, C2 – C8, inhibited respiration in accordance with their partition coefficients, suggesting that effects on mitochondrial respiration were related to membrane localization rather than cytosolic acidification. Supporting this, we present evidence that sorbic acid causes production of reactive oxygen species, the formation of petite (mitochondria-defective) cells, and Fe-S cluster defects. This work rationalises why yeasts that can grow in sorbic acid-preserved foods tend to be fermentative in nature. This may inform more-targeted approaches for tackling these spoilage organisms, particularly as the industry migrates to lower-sugar drinks, which could favour respiration over fermentation in many spoilage yeasts.

**IMPORTANCE:** Spoilage by yeasts and moulds is a major contributor to food and drink waste, which undermines food security. Weak acid preservatives like sorbic acid help to stop spoilage but some yeasts, commonly associated with spoilage, are resistant to sorbic acid. Different yeasts generate energy for growth by the processes of respiration and/or fermentation. Here we show that sorbic acid targets the process of respiration, so fermenting yeasts are more resistant. Fermentative yeasts are also those usually found in spoilage incidents. This insight helps to explain the spoilage of sorbic acid-preserved foods by yeasts and can inform new strategies for effective control. This is timely as sugar content of products like soft drinks is being lowered, which may favour respiration over fermentation in key spoilage yeasts.

## INTRODUCTION

Foods and beverages may be spoiled by yeasts and moulds despite the use of preservatives such as sorbic acid. Sorbic acid and acetic acid are both weak acids that inhibit many yeasts effectively, but weak acid resistant species can still cause contamination. Spoilage yeasts are a very small proportion of the overall numbers of known species. The number of recognised yeast species is in excess of 1500 (1) while those yeasts causing the majority of spoilage cases are limited to around 12 species (2). Similarly, only a small proportion of known mould species cause spoilage (3). Food lost to spoilage is a major food-security concern (4).

Complaints by consumers can be due to visible contamination, off-flavours or due to products being “blown”, or exploding in sealed packaging. This is caused by spoilage yeasts generating high levels of carbon dioxide through fermentation, and can result in physical injury (5). Yeast species found in factories producing foods or drinks have been characterized as belonging to Groups 1, 2 or 3, according to incidence and spoilage risk (6). *Z. bailii* is among a small number of spoilage yeasts categorised in Group 1. The Group 1 yeasts tend to be highly fermentative and show marked resistance to preservatives and osmotic stress. It is probable that most yeast species that have been recorded as fermentative (1) will also use respiration in different circumstances. In *S. cerevisiae*, high levels of glucose suppress respiration and the glucose is fermented (7). Respiration arises at lower glucose levels (≤ 0.5% - 1%, w/v). Furthermore, many yeast species are recorded as being non-fermentative, generating energy from carbon sources using respiration only. Approximately two thirds of all yeast species grow by respiration only (8). The Group 3 yeasts are very largely respiratory, and these species are sensitive to sorbic acid (6).

Many preservatives are weak acids, including propionic acid, sulphite (SO_2_), benzoic acid, and sorbic acid. It has previously been indicated that a major effect of these preservatives is acidification of the yeast cytoplasm (9-12). The weak acids enter cells rapidly by diffusion and can flow freely in-and-out (11, 13). As weak acids arrive in the yeast cytoplasm at neutral pH, they spontaneously dissociate to the anion, e.g., acetate, or bisulphite ion, and release H^+^. Large concentrations of weak acids can generate high levels of H^+^ and thereby lower the cytoplasmic pH, affecting protein structure and function (14). However, different weak acids inhibit yeasts at different levels; acetic acid requires ∼120mM to inhibit *S. cerevisiae*, while sorbic acid inhibits at ∼3mM. At these concentrations, acetic acid markedly lowers the internal pH (pH_i_) but sorbic acid only causes a very minor lowering of pH_i_ (11, 15). Therefore, the primary inhibition mechanism for sorbic acid and other similar weak acids (hexanoic acid, heptanoic acid, octanoic acid) has yet to be demonstrated. Partial inhibition of glycolysis rather than via pH_i_ has been suggested (15). Moreover, oil/water partition coefficients indicate that acetic acid should largely remain soluble in water whereas sorbic acid, which is more lipophilic, should largely occupy membranes. Certain membrane proteins have been shown to be specifically inhibited by sorbic acid (16, 17).

In the present study, based on a potential membrane action of sorbic acid and the observation that most sorbic-resistant spoilage yeasts grow by fermentation, we hypothesized, and then confirmed, that sorbic acid preferentially inhibits respiration over fermentation. A mechanistic explanation was then sought at the level of inhibition of mitochondrial function by sorbic acid. This insight should open a new avenue for understanding modes of weak-acid action and provide a new rationale for food preservation.

## RESULTS

### Sorbic acid sensitivity of respiratory growth in spoilage yeasts

We examined the effect of sorbic acid on growth of the model and food spoilage yeast, *S. cerevisiae*. As *S. cerevisiae* can decarboxylate sorbic acid, which requires the *PAD1* gene (18), a *S. cerevisiae* Δ*pad1* deletant (MIC 3mM sorbic acid, pH 4.0) was used for these growth experiments in order to prevent degradation of the weak acid during the experiment (such degradation enables later outgrowth, complicating discrimination of the normal growth phases). *S. cerevisiae* Δ*pad1* grew exponentially (∼2 hours doubling time) in YEPD pH 4.0, up to OD_600_ ∼10 (Fig. 1), and thereafter more slowly consistent with respiration of ethanol (Walker, 1998). Addition of 1 mM sorbic acid slowed the growth rate, to give a doubling time of ∼4.5 hours while 2 mM sorbic acid increased the doubling time to approximately 8.5 hours (Fig. 1). The growth yield at the end of the exponential growth phase was also decreased by the addition of sorbic acid, with an OD_600_ ∼7.5 achieved in 1 mM sorbic acid and OD_600_ ∼3.0 in 2 mM sorbic acid. At 2 mM sorbic acid, the subsequent, slow respiratory phase of growth appeared to be inhibited.

**FIG 1.**
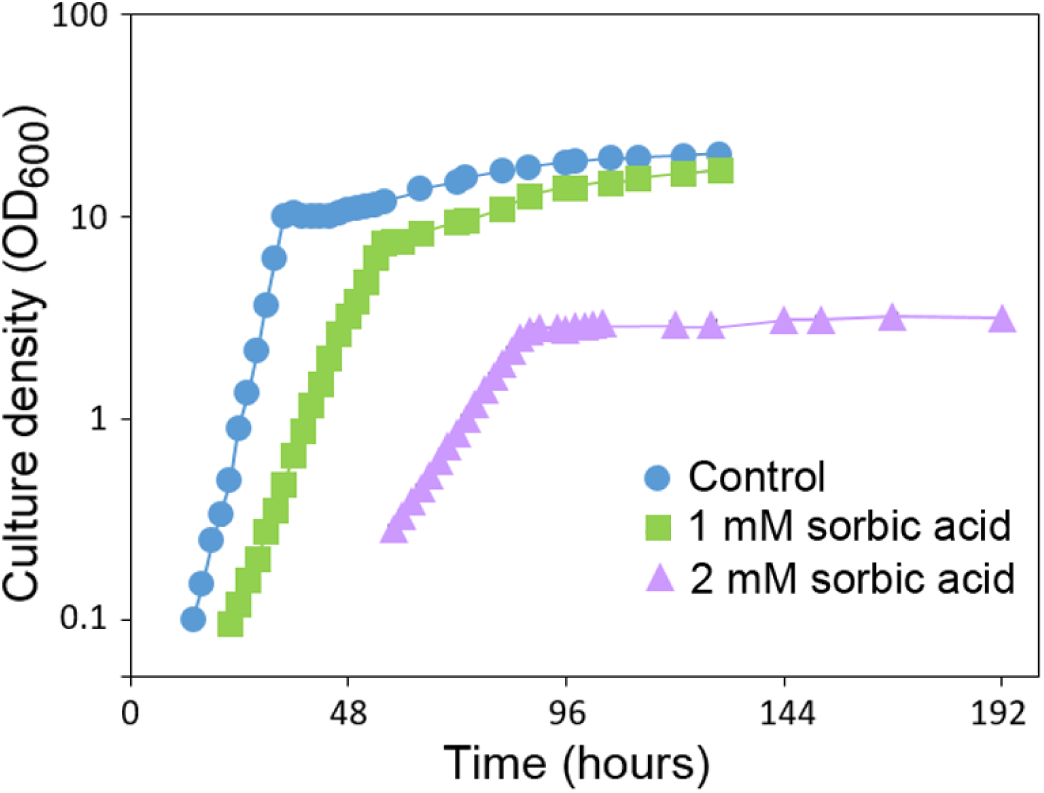
Effect of sorbic acid on growth of *S. cerevisiae* with glucose. *S. cerevisiae* Δ*pad1* was cultured with shaking at 120 rev. min^-1^, 24°C in flasks containing YEPD, pH 4.0 supplemented with the indicated concentrations of sorbic acid. Error bars (SD, n=3) were smaller than the dimensions of the symbols.

Since the respiratory phase of *S. cerevisiae* growth appeared to be stopped at 2 mM sorbic acid (Fig. 1), it was hypothesized that sorbic acid may selectively inhibit respiratory growth; so this was compared specifically by cultivation in YEP pH 4.0 supplemented with 30 g/l glycerol (no glucose). With 30 g/l glucose, growth was particularly inhibited at sorbic acid levels in excess of 2 mM sorbic acid, dropping to zero growth at the MIC of 3 mM sorbic acid (Fig. 2). In 30 g/l glycerol, full growth was limited to sorbic acid concentrations ≤0.7 mM and the MIC was 1.8mM sorbic acid. This indicated that respiratory growth of *S. cerevisiae* is hyper-sensitive to sorbic acid. A similar test was carried out with 30 g/l ethanol, an alternative respiratory substrate for *S. cerevisiae*, and again growth was inhibited at ∼1.5mM sorbic acid (data not shown).

**FIG 2.**
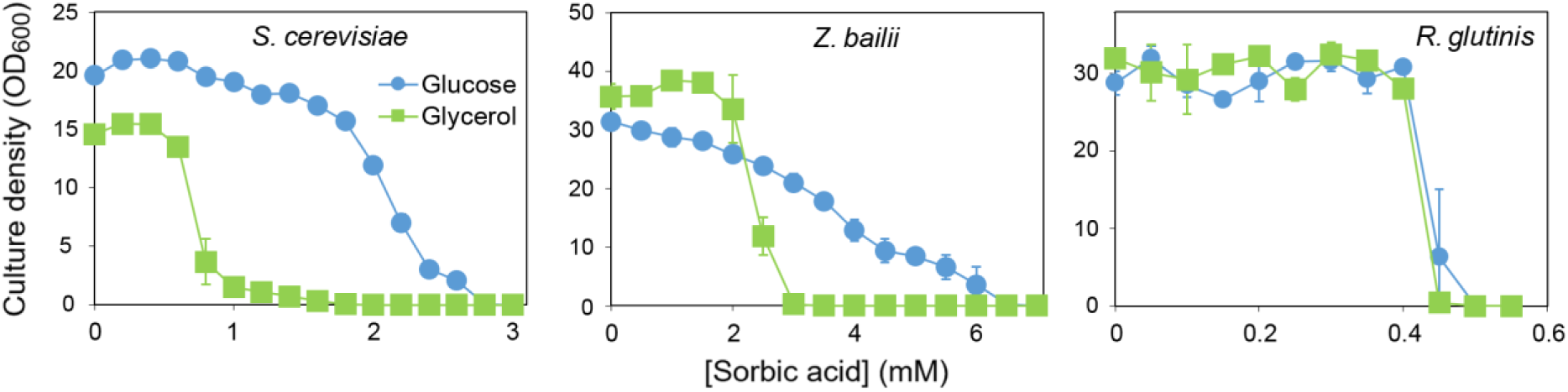
Growth inhibition by sorbic acid in yeasts cultured with glucose or glycerol. *S. cerevisiae, Z. bailii* or *R. glutinis* were cultured in either 3% (w/v) glucose or 3% glycerol, in YEP pH 4.0 supplemented with the indicated concentrations of sorbic acid. OD_600_ in flasks was determined after shaking at 120 rev. min^-1^, 24°C for 14 days. Points are means from three replicate determinations ± S.D.

Respiratory growth was also tested with the spoilage yeast *Z. bailii* in 30 g/l glycerol, and compared to growth in 30 g/l glucose (Fig. 2) (previously, *Z. bailii* has been reported to grow in glycerol, ethanol or acetic acid (19)). *Z. bailii* grew well in both 30 g/l glucose and in 30 g/l glycerol. However, the sorbic acid MIC in glycerol was 3.1 mM whereas in glucose the MIC was 6.6 mM sorbic acid. Therefore, both *Z. bailii* and *S. cerevisiae* were inhibited at lower levels of sorbic acid when growing by respiration.

A similar experiment was carried out with *Rhodotorula glutinis*, which has been reported as fermentation deficient in glucose and other sugars (1, 8). This red yeast grew similarly well either in 30 g/l glucose or 30 g/l glycerol in the shaking flasks (Fig. 2). However, *R. glutinis* was hyper-sensitive to sorbic acid, and the MIC was almost identical at ∼0.5 mM acid whether in 30 g/l glucose or 30 g/l glycerol. As sorbic acid resistance in glucose (seen with *S. cerevisiae* or *Z. bailii*) was absent in this respiration-only yeast, the results further corroborate that sorbic acid selectively inhibits respiratory growth.

### Relationship between sorbic acid sensitivity and respiratory or fermentative growth of diverse yeast species

Above, *S. cerevisiae, Z. bailii* and *R. glutinis* had all been grown in 30 g/l glucose or 30 g/l glycerol during treatment with sorbic acid. To ascertain whether the key observations with these species extended to other yeasts, a further 14 species were tested in a similar way, as representatives of Davenport Groups 1, 2, and 3 (Table 1). On glucose, as expected Group 1 yeasts were highly resistant to sorbic acid, Group 2 were moderately resistant, and Group 3 were sensitive to sorbic acid (Table 1). All four species in Group 3 exhibited similar sensitivity to sorbic acid whether in 30 g/l glucose or 30 g/l glycerol. As indicated above for *R. glutinis*, these species were all fermentation-defective in glucose (Table S3) so can be assumed to be respiring in both glucose and glycerol. Within Group 2, three of the test species show moderate fermentation (*Wicherhamomyces anomalus, Candida pseudointermedia, Candida parapsilosis*) while two have high fermentation (*Saccharomyces cerevisiae* and *Torulaspora delbruckii*) (Table S3). The MICs of sorbic acid for these yeasts in the respiratory substrate glycerol were ∼60%–90% of the MICs observed in glucose (Table 1). This indicated that fermentative metabolism by the Group 2 yeasts was associated with a moderate elevation of sorbic acid resistance. Sorbic acid resistance of the high-fermentation Group 1 yeasts tended to be the most strongly affected by carbon source. The sorbic acid MIC for these species when growing by respiration was between 40% - 65% of their MICs when growing by fermentation.

**TABLE 1.**
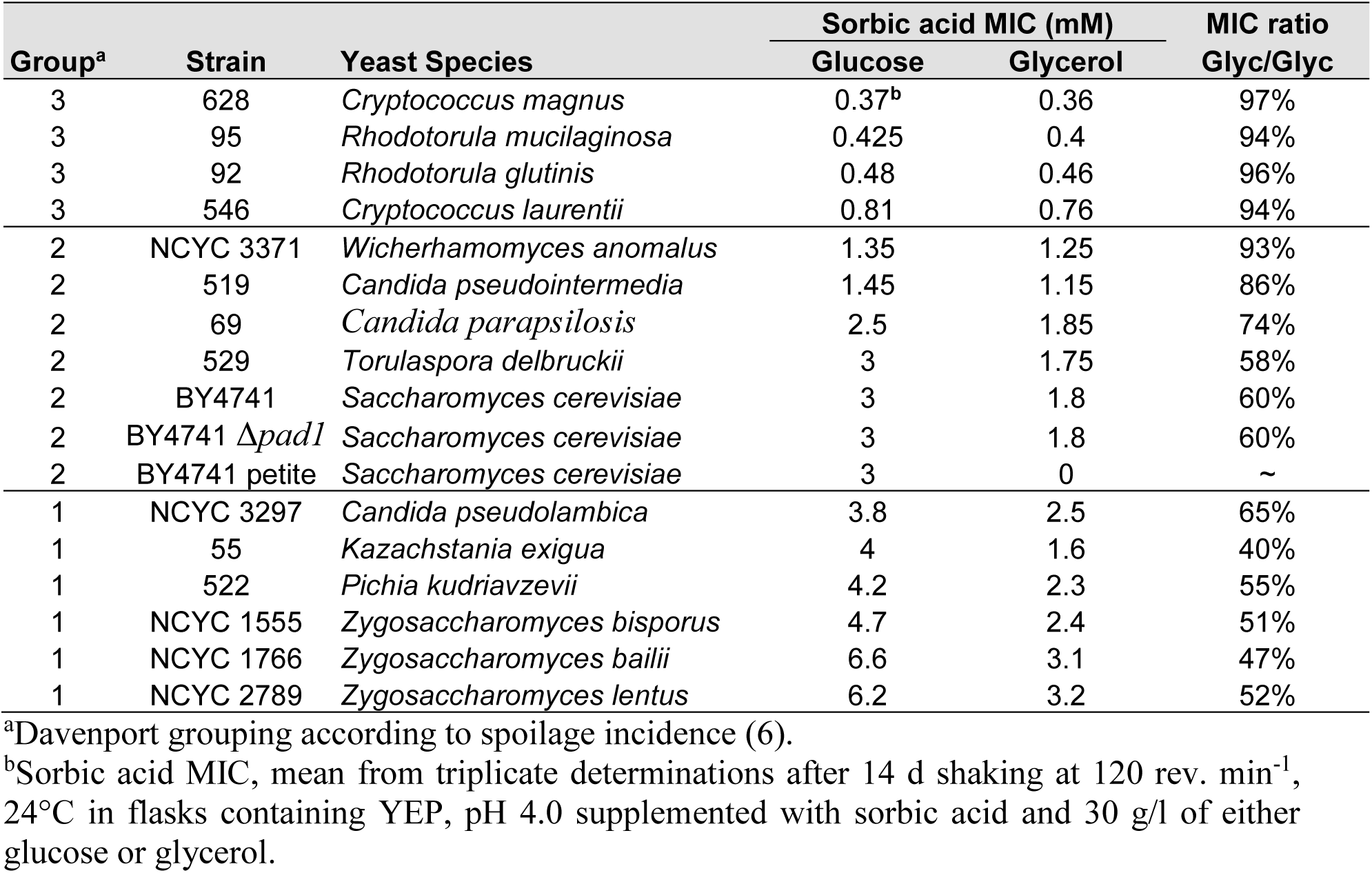
Respiratory growth sensitizes spoilage species to sorbic acid.

In glucose, there was overlap in the fermentation rates of the Group 1 and 2 yeasts (Table S3) but not in their MICs (Table 1). Therefore, a high level of fermentation alone did not appear to be sufficient to explain the highest levels of sorbic acid resistance. To test more rigorously any relationship between fermentation activity and sorbic acid resistance, we tested 687 yeast strains, representing 191 yeast species (Table S2), for sorbic acid MICs and fermentation level in 180 g/l glucose (this higher glucose level accentuates differences in fermentation capacity, Table S3). The data for different strains of each yeast species were averaged before plotting. There was a weak but significant positive correlation between fermentation and sorbic acid resistance (correlation, R^2^ = 0.3058; p < 0.0001) (Fig. 3). The bulk of the 53 species that showed no fermentation were found to be sensitive to sorbic acid, with just a few showing moderate resistance (e.g. *Yarrowia lipolytica*). The species with the greatest resistance tended to be those with the highest fermentation.

**FIG 3.**
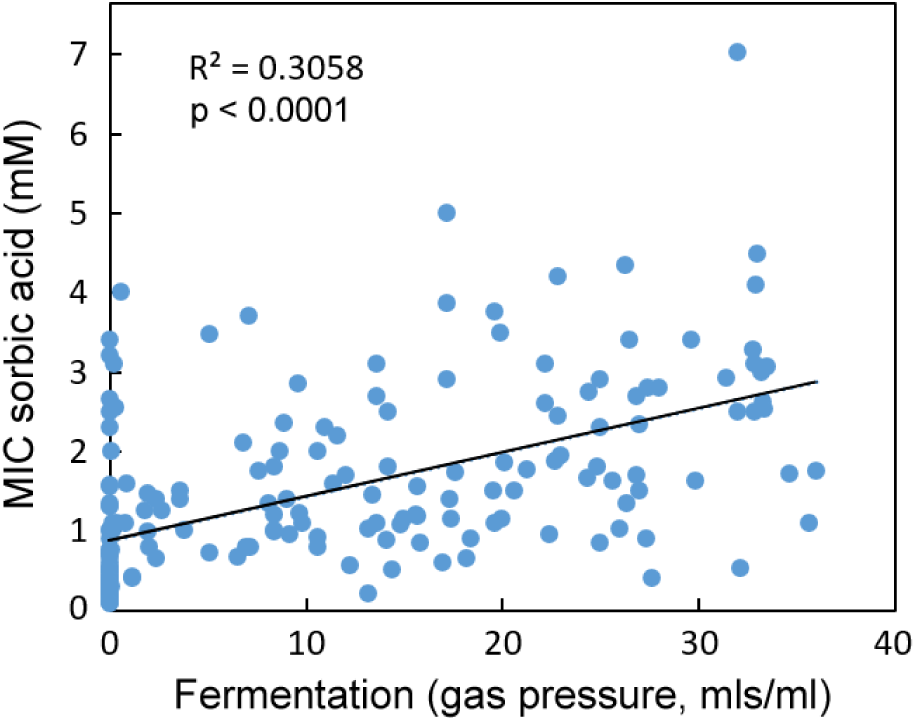
Comparison of the level of fermentation and sorbic acid resistance exhibited by 191 yeast species (encompassing 687 yeast strains). Organisms are listed in Table S2. Sorbic acid resistance (MIC) was determined in YEPD (pH 4.0) after 14 days at 25^°^C, and the fermentation was tested after 28 days in static bottles in YEP with 180g/l (1 M) glucose. Fifty three species showed zero fermentation. The slope was fitted by linear regression. R^2^ and p values (shown on the figure) were determined by Pearson correlation.

### Selective inhibition of respiratory activity by sorbic acid in yeasts

As relative dependence on respiration for growth should increase with decreasing fermentation, and respiration-only species were the most sorbic acid sensitive, we reasoned that respiratory metabolism could be a target of sorbic acid. About two-thirds of yeast species are recorded as non-fermentative (8), so considered respiration only. We tested whether respiration is inhibited by sorbic acid using *S. cerevisiae* growing in 30 g/l glycerol, using Warburg manometry. The data substantiated that the yeast was respiratory in these conditions, absorbing oxygen and producing equivalent carbon dioxide (Fig. 4). Inclusion of sorbic acid at the MIC level (1.8 mM) strongly inhibited respiration. After 120 min, oxygen removal and carbon dioxide production in the presence of sorbic acid were ∼20% of the control level. In the case of the major spoilage yeast *Z. bailii*, respiratory growth in 30 g/l glycerol was associated with slightly higher oxygen absorption than carbon dioxide production, but these parameters were both reduced to a similar level at 3.5 mM sorbic acid (approximating the relevant MIC for this organism). The experiment was repeated with the respiration-only yeast *Rhodotorula glutinis* in glycerol (Fig. 4). Sorbic acid at the relevant MIC (0.46 mM) inhibited the respiration of *R. glutinis*, to a similar extent as observed in *S. cerevisiae* and *Z. bailii*.

**FIG 4.**
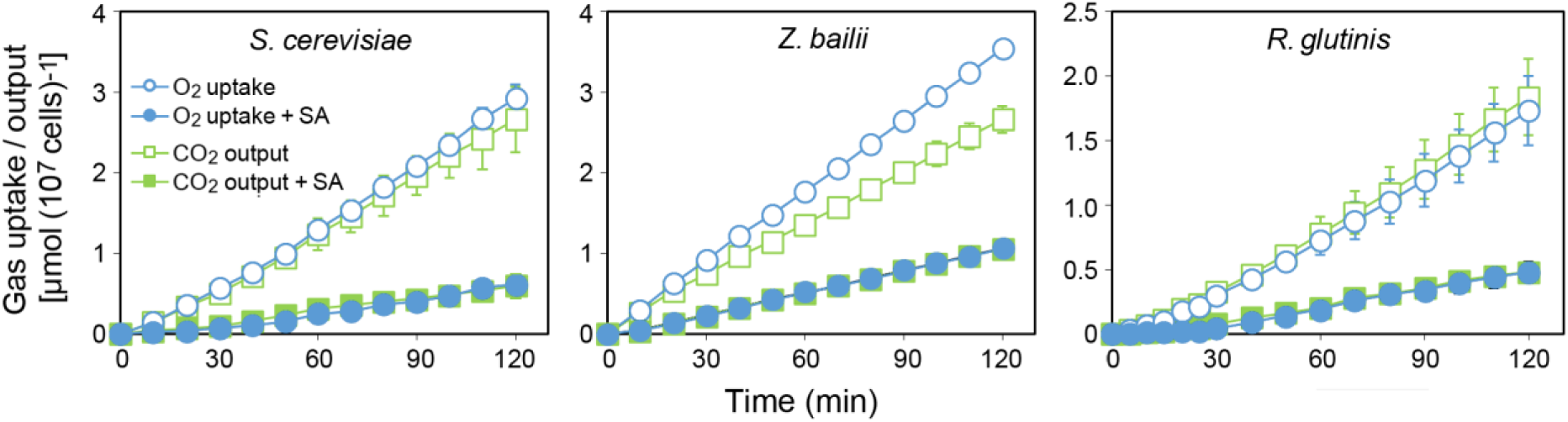
Inhibition of respiration by sorbic acid. Yeasts growing with shaking at 24°C in YEP pH 4.0 containing 3% (w/v) glycerol were monitored for O_2_ uptake and CO_2_ output by Warburg manometry. Where indicated (closed symbols) sorbic acid (SA) was included at 1.8 mM for *S. cerevisiae*, 3.5 mM for *Z. bailii* or 0.46 mM for *R. glutinis*. Points are means from three replicate determinations ± S.D.

The relative sorbic acid sensitivities of respiration and fermentation were compared in *S. cerevisiae*. During culture in glycerol and exposure to sorbic acid (1.6 mM), respiration was inhibited by ∼68% and did not recover over longer treatment times up to 6 hours (Fig. 5). The fermentation rate during culture with 2% glucose was less strongly inhibited, by ∼48%, immediately following sorbic acid addition. Furthermore, fermentation quickly recovered to pre-treatment levels after 2.5 hours exposure to sorbic acid. This comparison was further tested at 0.5% glucose, a concentration that allows both fermentation and respiration in *S. cerevisiae* BY4741. Again, sorbic acid inhibited respiration by ∼60% and this did not recover after 3 hours, whereas there was less (∼38%) inhibition of fermentation, which largely recovered after 3 hours (Fig. 5). The results indicate that respiration is more sensitive than fermentation to inhibition by sorbic acid.

**FIG 5.**
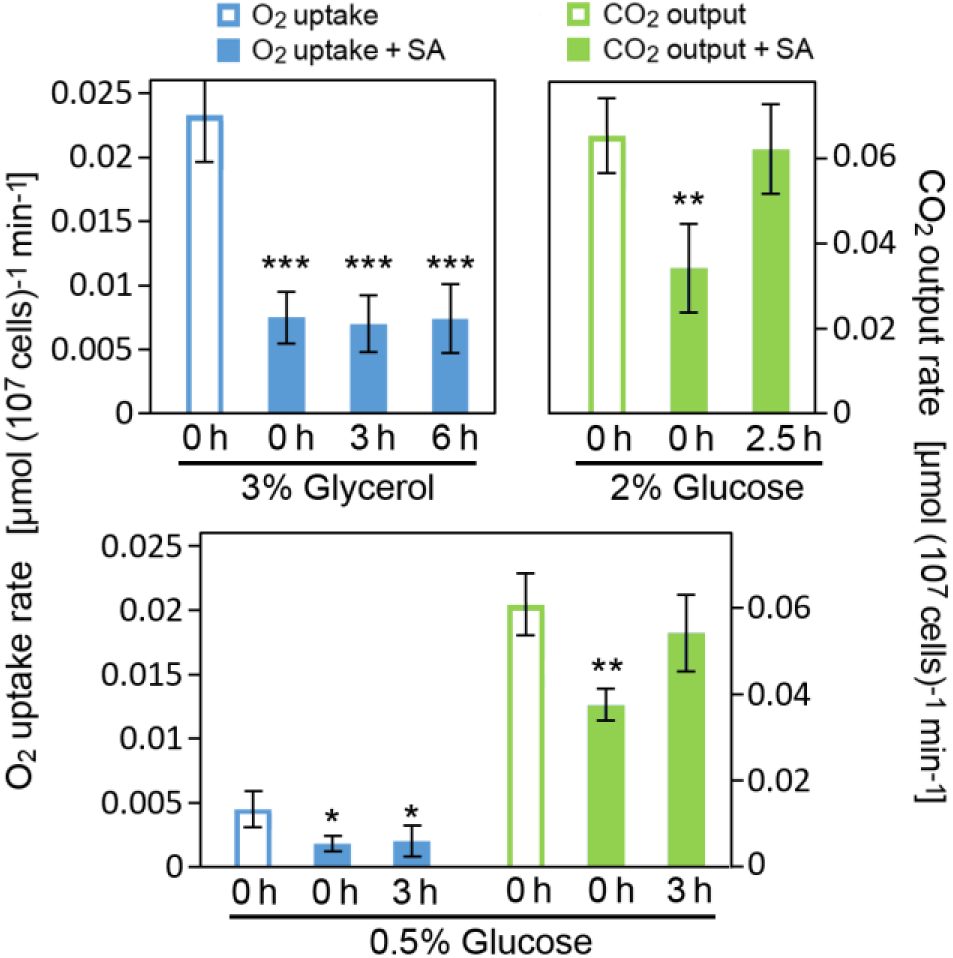
Comparative **i**nhibition of respiration and fermentation by sorbic acid. *S. cerevisiae* growing with shaking at 24°C in YEP pH 4.0 containing either glucose or glycerol at the indicated concentrations (w/v) was either treated or not with 1.6 mM sorbic acid (SA) and monitored for O_2_ uptake and CO_2_ output. Measurements were made for up to 40 min either immediately following sorbic acid treatment (0 h) or after the later exposure-times indicated. Mean data are shown from four replicate determinations ± S.D. *p<0.05, **p<0.01, ***p<0.001, according to Student’s t test, two tailed

### Selective inhibition of respiratory growth by different weak acids is correlated with chain length and membrane solubility

We also examined acetic acid, as a weak acid that is more hydrophilic (less membrane soluble) than sorbic acid. Much higher concentrations of acetic acid were required to inhibit yeast growth than sorbic acid. Moreover, unlike with sorbic acid, inhibition of *S. cerevisiae* by acetic acid was similar in 30g/l glycerol or 30g/l glucose, with MICs close to 140 mM (Fig. S1). This outcome was reflected also with *Z. bailii*, which was highly resistant to acetic acid in glycerol and in glucose, where the MIC levels were similar at ∼450 mM acetic acid (Fig S1). To explore further potential relationships between weak acid-sensitivity of respiratory growth and weak acid hydrophobicity, a wider range of weak acids with two- to eight-carbon lengths was tested in *S. cerevisiae*. MICs were determined for each weak acid during growth in both glycerol and glucose; the ratio between these MIC values provided an indication of the relative sensitivity of respiratory growth to each weak acid. In both media, the MICs declined markedly with increasing carbon chain length of the weak acids (Table S4). However, the relative decline in MIC differed in glycerol versus glucose. As a result, there was a tight inverse relationship between carbon chain length and relative inhibition of respiratory versus fermentative growth by the weak acids (Fig. 6). The longer-chain acids have a relatively high octanol/water partition coefficient (cLogP), predictive of greater membrane localization. Acetic acid is relatively hydrophilic and gave similar MICs during respiration (in glycerol) or fermentation (glucose); whereas, the MICs of 6C (hexanoic) and 8C (octanoic) acids in glycerol were only 68% and 53% of those in glucose.

**FIG 6.**
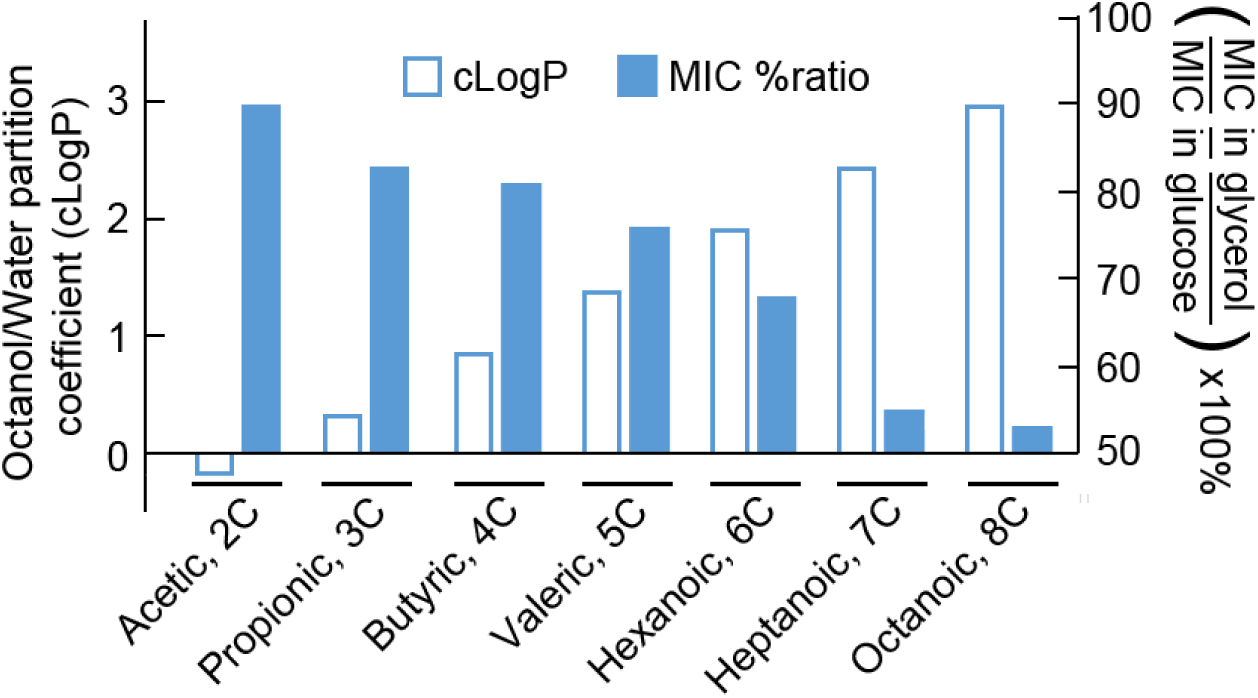
Membrane-partitioning tendency of acids correlates with relative toxicities to respiring *S. cerevisiae*. MIC, minimum inhibitory concentration. Determined after 14 d shaking at 120 rev. min^-1^, 24°C, in flasks containing YEP, pH 4.0 supplemented with weak acid and 30 g/l of either glucose or glycerol. Underlying values are given in Table S4.

### Mechanisms underlying sorbic acid-sensitivity of yeast respiration

The above data collectively support the hypothesis that longer chain-length weak acids, like sorbic acid, selectively target respiration by yeasts, and that this effect is correlated with membrane solubility. Membrane perturbation and effects on the respiratory chain are commonly associated with production of reactive oxygen species (ROS) (20, 21). In the present study, analysis using the ROS probe DHE indicated that sorbic acid, at sub-inhibitory concentrations, promotes ROS production in *S. cerevisiae* (Fig. 7A). One consequence of mitochondrial ROS production in organisms like *S. cerevisiae* is the formation of petite (mitochondria-defective) cells, arising from mitochondrial DNA damage; mitochondrial DNA encodes mainly for proteins that are part of the respiratory chain (22). We tested whether petite-cell formation could be a contributor to the apparent selective targeting of respiration during sorbic acid treatment. We grew *S. cerevisiae* on YEPD agar supplemented or not with sorbic acid, then assayed for respiratory competency by replica plating colonies to YEP-glycerol agar. There was a ∼2.3-fold increase in petite-cell frequency in the presence of 0.75 mM sorbic acid compared to control (Fig. 7B). ROS are also known to impair iron-sulfur cluster (ISC) biogenesis, which takes place in the mitochondria. We tested whether mutants of the ISC biogenesis pathway were hypersensitive to sorbic acid treatment. On glycerol, the mutants *bol3Δ* and *nfu1Δ* and to some extent *isu1Δ* were more sensitive to the weak acid than the wild type strain, suggesting an effect of sorbic acid on iron-sulfur (FeS) protein formation (Fig. 7C). Isu1 is a scaffold protein involved in the first step of [2Fe-2S] cluster assembly while Nfu1, assisted by Bol3, facilitates the transfer of [4Fe-4S] clusters from the assembly complex to client proteins while protecting the co-factor from oxidative damage (23) (Fig. 7C). Succinate dehydrogenase, essential for respiration, is one of these client proteins. The collective data suggest that sorbic acid generates ROS in mitochondria, which could lead to depletion of functional respiratory complexes through pathways including petite cell formation and FeS cluster targeting.

**FIG 7.**
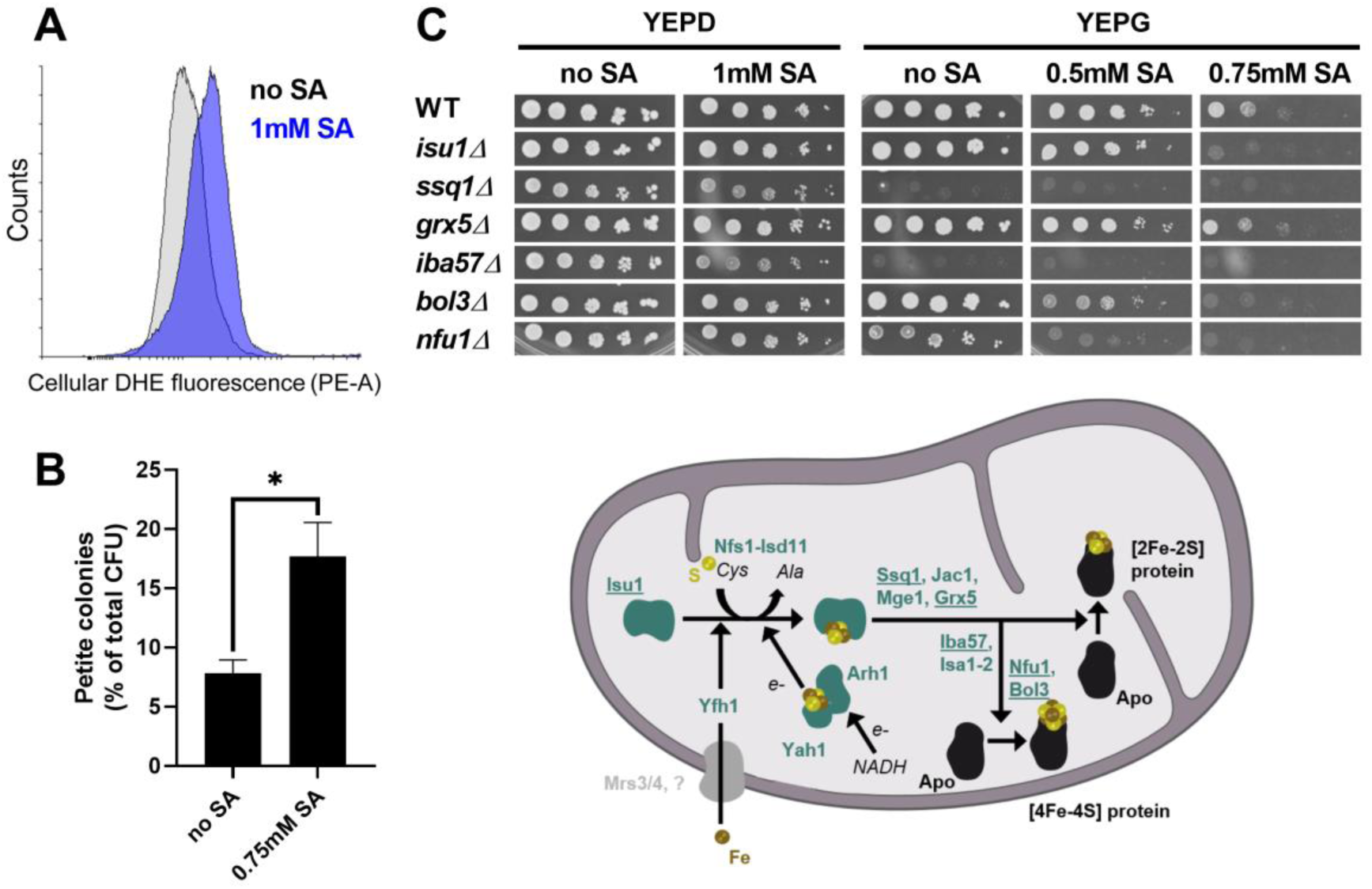
ROS production and damage to mitochondrial function with sorbic acid during respiration. (A) ROS production upon sorbic acid treatment was measured using the fluorescent probe DHE after 4 h incubation of *S. cerevisiae* with sorbic acid. (B) *S. cerevisiae* colonies cultivated for at least 7 days with or without 0.75 mM sorbic acid (i.e., sub-inhibitory concentration) were replica-plated onto YEP-glucose (YEPD) and YEP-glycerol (YEPG) to assess petite formation (petite cells do not grow on YEPG). Mean data are shown from triplicate independent growth experiments ± SEM. *p < 0.05 according to Student’s t test, two tailed. (C) Upper panel: Serial dilutions of *S. cerevisiae* BY4743 and the indicated isogenic deletion strains were spotted onto agar plates containing a fermentable (YEPD) or respiratory (YEPG) carbon source with or without sorbic acid and incubated at 30°C for 2 or 4 days, respectively. Lower panel: simplified scheme showing proteins involved in the biogenesis of FeS-clusters and their transfer to mitochondrial apo-proteins.

## DISCUSSION

This study shows that respiration by yeasts is more sensitive than fermentation to inhibition by the major food preservative sorbic acid. Furthermore, fermentative metabolism shows greater recovery than respiratory metabolism over time after sorbic acid shock. This is an important result for industry and helps explain why the highly fermentative species such as *Z. bailii* can cause such catastrophic spoilage incidents. Indeed, we show that historical categorization of yeast species according to their propensity to cause food or beverage spoilage (6) maps closely to their capacities to ferment or respire, with the yeasts least associated with spoilage (Group 3) being fermentation-defective and most reliant on respiration.

Respiration in glycerol was shown to be inhibited by sorbic acid in *S. cerevisiae* and 14 other yeast species showed sorbic acid-sensitive respiratory growth. In addition, a screen of 191 yeast species established a correlation between fermentative activity and sorbic acid resistance. Previous tests carried out on a mould, *Aspergillus niger*, had shown that sorbic acid inhibited respiration and germination in asexual spores (24). However, fermentation is low or absent in such filamentous fungi, precluding the type of comparisons established here with yeasts. Respiratory growth was inhibited by sorbic acid at 3.1mM for *Z. bailii* (Group 1), 1.8 mM for *S. cerevisiae* (Group 2) and 0.46 mM for *R. glutinis* (Group 3), but these relative differences were accentuated in glucose, where growth can occur by fermentation in *Z. bailii* and *S. cerevisiae*.

Other weak acids also inhibited respiratory growth in *S. cerevisiae* (Table S4), and the toxic effects of the different weak acids was much greater with the longer-chain weak acids. This strongly suggests that the relative toxic effect is related to lipophilicity and the weak acid being absorbed into lipid membranes. A weak acid such as decanoic acid (10-carbons) is absorbed so profusely into the membranes that the membranes burst, causing rapid cell death (25). Previous studies have indicated that yeast respiration can be inhibited by certain weak acids (26-28). However, such earlier work typically has not distinguished whether these are selective (causative) effects, targeting respiration, as opposed to broader inhibition of cell activities by sorbic acid among which decreased respiration would be just one of several effects. The ability to dissect respiratory from fermentative growth in yeasts provided a valuable tool to resolve these possibilities in this study.

How does sorbic acid target respiration? To help address this question, we initially had attempted a screen of the yeast homozygous deletant collection (29, 30) to find gene functions required for respiratory sensitivity to sorbic acid. Although a number of deletants on glycerol did exhibit higher sorbic acid resistance than the wild type, particularly an *atg9* deletant defective in autophagy, none of these identified gene functions were involved in respiration (I. Geoghegan and S.V. Avery, Unpublished data). Instead, we considered the correlation between the respiratory inhibition and membrane solubilities of the different weak acids (above), and the ROS production that is commonly associated with mitochondrial-membrane perturbation (20, 21). Our results showed that sorbic acid can promote ROS production. Furthermore, key effects of ROS on cellular function (31-33) could be seen in sorbic acid treated cells, i.e., formation of mitochondria-defective petite cells and indications of FeS-cluster pathway defects. The mitochondrial DNA damage that typically produces petite cells is normally irreversible. Therefore, the increased incidence of petite cells could help to explain the non-recovery of respiratory activity that we observed following sorbic acid treatment, whereas fermentation (which can continue in petite cells) did recover. It was also notable that specific FeS-pathway functions which conferred sorbic acid resistance are involved in shielding FeS delivery from ROS, to client proteins like succinate dehydrogenase which is essential for respiration. We propose a model in which ROS generated by membrane-localized sorbic acid causes depletion of mitochondrial respiratory function, through pathways including petite cell formation and FeS cluster targeting.

It has been argued that population heterogeneity (preservative hetero-resistance) among individual cells or spores within spoilage-yeast or -mould populations has major implications for food spoilage (34-37). Accordingly, it may take only a few, preservative hyper-resistant cells to initiate spoilage. It was previously reported that *Z. bailii* causes spoilage at a level of contamination of one cell/bottle (38). Although heterogeneity was not a focus of this study, our finding that mitochondrial function is a key determinant of sorbic acid sensitivity is notable in this context. That is because mitochondrial activity and cellular redox status are quite heterogeneous within cell populations (39, 40). In the present study, petite cell formation with sorbic acid was not uniform across the yeast cell populations. Therefore, it is possible that mitochondrial heterogeneity could be one factor determining the preservative hetero-resistance of spoilage yeasts, the mechanistic bases for which have yet to be resolved.

Overall, this paper shows that respiration is selectively inhibited by sorbic acid, seemingly in a wide range of yeast species. We propose that ROS generated by membrane-localized weak acid could explain this respiratory targeting. These findings are especially timely as the beverages industry seeks to reduce sugar content of its products, in response to soft-drinks sugar taxes introduced by a number of governments. Because of catabolite repression, as glucose is decreased, respiratory activity of spoilage yeasts increases (assuming there is available oxygen) (7). Our results suggest that an increased reliance of spoilage yeasts on respiration for growth in lower-sugar beverages is likely to strengthen the preservative efficacy of weak acids. In contrast, there is evidence that sugar-substituted foods can be more prone to spoilage by moulds (41), organisms that are more reliant on respiration regardless of glucose concentration. It seems that the profile of organisms associated with spoilage incidents is likely to change as the glucose content of soft drinks is lowered.

## MATERIALS AND METHODS

### Yeast species and strains

The principal yeast species and strains used in this study are listed in Table S1, which records both previous (8) and updated (1) strain numbers and species names, and the sources of the yeasts. In addition, 687 yeast strains spanning 191 species were tested for the relationship between fermentation and sorbic acid resistance, and these are listed in Table S2. The identities of all the strains were determined by sequencing the D1/D2 region of the 26S rDNA (42). Experiments with *S. cerevisiae* were with strain BY4741 unless specified otherwise. Yeasts were stored in glycerol on ceramic beads at – 80°C (Microbank™), and maintained in the short term on MEA (malt extract agar, Oxoid) slopes at 4°C.

### Growth conditions

The growth medium for routine culturing was YEPD, containing 20g/l glucose, 20g/l bacteriological peptone (Oxoid), and 10g/l yeast extract (Oxoid). The medium was adjusted to pH 4.0 with 5M HCl prior to sterilization by autoclaving. Where specified, the glucose concentration was amended to 5g/l, 10g/l, 30g/l, or 180g/l. Other experiments required yeast extract and peptone, YEP, with 30g/l glycerol adjusted to pH 4.0. Some batches of peptone or yeast extract contained low levels of glucose, so these were avoided when zero or low levels of known amounts of glucose were required.

Starter cultures were grown in 10ml YEPD pH 4.0 in 28ml McCartney bottles, inoculated with yeast from MEA slopes. Bottles were incubated statically for 48 hours at 24°C. These starter cultures were used to inoculate experimental cultures, which comprised either 5ml or 10ml YEPD, pH 4.0 in 28ml McCartney bottles, or 40ml YEPD, pH 4.0 in 100ml conical flasks which were shaken at 120 rpm, 24°C.

Sorbic acid was dissolved in methanol to make stock solutions of 100mM, 200mM or 400mM. Aliquots from the stock solution appropriate for the desired final sorbic-acid concentration were transferred to the medium before the pH 4.0 adjustment. At pH 4.0, ∼ 85% of added sorbic acid exists as free acid and ∼15% as anion. Acetic acid and propionic acids (liquids) were added directly to media before adjusting the pH to 4.0. Butyric acid, valeric acid, hexanoic acid, heptanoic acid and octanoic acid (liquids) were diluted in methanol. The final methanol levels in media did not exceed 1% (v/v) (20% - 30% methanol is required to exert toxicity in yeast).

### Measurement of gas pressure from fermentation

Fermentation by yeasts uses sugars, commonly glucose, to generate ATP accompanied by metabolism of sugar to ethanol and CO_2_ (1, 8). The level of fermentation was estimated by measuring the gas pressure generated by the yeast. Yeasts were inoculated into 10ml YEPD pH 4.0, in sealed triplicate 28ml McCartney bottles, and incubated at 24°C for up to 28 days. Glucose was routinely used at 20g/l, but 5g/l, 10g/l and 30g/l were also tested. Pressure in the sealed bottles was tested at 28 days with a gas syringe (average gas volumes were between 0 – 30ml, when adjusted to atmospheric pressure). A parallel fermentation was carried out using 180g/l glucose (1M), in 5ml YEPD pH 4.0 in the sealed bottles. Gas volumes up to 200ml were obtained in the 28ml McCartney bottles (approx. 10 atmospheres). Gas pressure was calculated as the volume (mls) of gas generated per 1ml of medium.

### Warburg manometry

A Warburg manometer was used to measure the absorption of O_2_ and the efflux of CO_2_ from yeast, by determining gas pressures over a period of 70 min. The shaking flasks contained 3 ml of medium and either 0.4ml of water or 20% KOH held at 24°C. For measuring respiration, glucose-free YEP pH 4.0 medium was supplemented with glycerol at 30g/l and yeasts were pre-cultivated in this medium for 12-14 hours at 24°C then transferred to fresh medium at 10^7^ cells/ml prior to measurements in the manometer. Respiration was confirmed as the sole metabolic route where absorption of oxygen and efflux of carbon dioxide were almost identical. Fermentation (carbon dioxide efflux but no absorption of oxygen) was tested in medium supplemented with glucose (20 g/l) rather than glycerol, after pre-cultivation in this medium for 12-14 hours at 24°C before transfer to fresh medium at 10^7^ cells/ml. A lower glucose concentration (5 g/l) was used to assess both respiration and fermentation.

### ROS accumulation

Detection of cellular reactive oxygen species (ROS) was with the fluorescent probe dihydroethidium (DHE) (43). Samples of yeast culture (*S. cerevisiae* BY4743) grown in YEPD, (pH 4.0) with or without sorbic acid for 4 hours were centrifuged, washed and then incubated in 100 µl PBS with 5 µM DHE for 30 min at 30°C, 120 rev.min^-1^. Cells were then harvested by centrifugation and resuspended in 500 µl PBS before analysis of cellular DHE fluorescence using a Beckman Astrios MoFlo cell sorter equipped with a 488 nm laser.

### Petite cell formation

Yeast cells were spread plated and grown for at least 7 days on YEPD agar, supplemented or not with 0.75 mM sorbic acid, then colonies were replicated to fermentable (YEPD) or respiratory (YPG) solid medium. After 3 days, the percentage of petite colonies was assessed according to respiratory deficiency (no growth) on YPG versus YEPD. For the sorbic acid supplemented YEPD (above), as low-pH agar degenerates with heating and will not subsequently set, agar medium at near-neutral pH was autoclaved before acidifying to pH 4.0 just before pouring. The YEPD was prepared without agar (not pH-adjusted) and sorbic acid added to defined concentrations. Samples (50 ml) were removed from each medium batch that contained sorbic acid and titrated to pH 4.0 with 5M HCl to determine the volume of acid needed to adjust each batch to pH 4.0. Agar was added (16g/l) to the neutral non-pH adjusted medium, which was then warmed to melt the agar before autoclaving. During subsequent cooling, the medium was held at 50°C, before acidification to pH 4.0 with the appropriate, pre-determined volume of acid, and poured into Petri dishes. The agar had no effect on pH or buffering.

## ACKNOWLEDGEMENTS

This work was supported by the Biotechnology and Biological Sciences Research Council [grant number BB/N017129/1]. This is an Industry Partnering Award in conjunction with Lucozade Ribena Suntory and Mologic Ltd.

## LEGENDS TO SUPPLEMENTARY FIGURES (data provided in separate attachments)

**TABLE S1** Principal yeast species used in this research. Original and current species names are provided (1) together with the strain sources.

**TABLE S2** Full list of yeast species, encompassing 687 strains, which were tested for fermentation and sorbic acid MIC (results summarised in Fig. 3).

**TABLE S3** Fermentation by spoilage and non-spoilage yeast species.

**TABLE S4** Resistance of *S. cerevisiae* to weak acids with different carbon chain lengths.

**FIG S1** Growth on glucose or glycerol in the presence of acetic acid. *S. cerevisiae* (top panel) or *Z. bailii* (bottom panel) were cultured in either 30g/l glucose (open squares) or 30g/l glycerol (closed squares), in YEP pH 4.0 supplemented with the indicated concentrations of acetic acid. OD_600_ in flasks was determined after shaking at 120 rev. min^-1^, 24°C for 14 days. Points are means from three replicate determinations ± S.D.

## REFERENCES

1. Kurtzman CP, Fell JW, Boekhout T. 2011. The Yeasts, a Taxonomic Study 5th ed. Elsevier, Amsterdam.

2. Stratford M, James SA. 2003. Non-alcoholic beverages and yeasts, p 309–345. In Boekhout T, Robert V (ed), Yeasts in Food Beneficial and Detrimental Aspects. Behr’s-Verlag, Hamburg.

3. Pitt JI, Hocking AD. 2009. Fungi and Food Spoilage, Third Edition, 3rd ed. Springer, New York.

4. Avery SV, Singleton I, Magan N, Goldman GH. 2019. The fungal threat to global food security. Fungal Biol 123:155–157.

5. Grinbaum A, Ashkenazi I, Treister G, Goldschmiedreouven A, Block CS. 1994. Exploding bottles - eye injury due to yeast fermentation of an uncarbonated soft drink. Brit J Ophthalmol 78:883.

6. Davenport RR. 1998. Microbiology of soft drinks, p 197-216. In Ashurst PR (ed), The Chemistry and Technology of Soft Drinks and Fruit Juices. Sheffield Academic Press, Sheffield, UK.

7. Walker GM. 1998. Yeast Physiology and Biotechnology. Wiley, UK.

8. Barnett JA, Payne RW, Yarrow D. 2000. Yeasts: Characteristics and Identification, 3rd ed. Cambridge University Press, Cambridge, UK.

9. Fernandez-Nino M, Marquina M, Swinnen S, Rodriguez-Porrata B, Nevoigt E, Arino J. 2015. The cytosolic pH of Individual *Saccharomyces cerevisiae* cells is a key factor in acetic acid tolerance. Appl Environ Microbiol 81:7813–7821.

10. Orij R, Postmus J, Ter Beek A, Brul S, Smits GJ. 2009. In vivo measurement of cytosolic and mitochondrial pH using a pH-sensitive GFP derivative in *Saccharomyces cerevisiae* reveals a relation between intracellular pH and growth. Microbiology-SGM 155:268–278.

11. Stratford M, Nebe-von-Caron G, Steels H, Novodvorska M, Ueckert J, Archer DB. 2013. Weak-acid preservatives: pH and proton movements in the yeast *Saccharomyces cerevisiae*. Int J Food Microbiol 161:164–171.

12. Ullah A, Chandrasekaran G, Brul S, Smits GJ. 2013. Yeast adaptation to weak acids prevents futile energy expenditure. Front Microbiol 4.

13. Palma M, Guerreiro JF, Sa-Correia I. 2018. Adaptive response and tolerance to acetic acid in *Saccharomyces cerevisiae* and *Zygosaccharomyces bailii*: A physiological genomics perspective. Front Microbiol 9:274.

14. Krebs HA, Wiggins D, Stubbs M, Sols A, Bedoya F. 1983. Studies on the mechanism of the anti-fungal action of benzoate. Biochem J 214:657–663.

15. Burlini N, Pellegrini R, Facheris P, Tortora P, Guerritore A. 1993. Metabolic effects of benzoate and sorbate in the yeast *Saccharomyces cerevisiae* at neutral pH. Arch Microbiol 159:220–224.

16. Bauer BE, Rossington D, Mollapour M, Mamnun Y, Kuchler K, Piper PW. 2003. Weak organic acid stress inhibits aromatic amino acid uptake by yeast, causing a strong influence of amino acid auxotrophies on the phenotypes of membrane transporter mutants. Eur J Biochem 270:3189–3195.

17. Melin P, Stratford M, Plumridge A, Archer DB. 2008. Auxotrophy for undine increases the sensitivity of *Aspergillus niger* to weak-acid preservatives. Microbiology-SGM 154:1251–1257.

18. Stratford M, Plumridge A, Archer DB. 2007. Decarboxylation of sorbic acid by spoilage yeasts is associated with the *PAD1* gene. Appl Environ Microbiol 73:6534–6542.

19. Sousa MJ, Miranda L, CorteReal M, Leao C. 1996. Transport of acetic acid in *Zygosaccharomyces bailii*: Effects of ethanol and their implications on the resistance of the yeast to acidic environments. Appl Environ Microbiol 62:3152–3157.

20. Fedoseeva IV, Pyatrikas DV, Stepanov AV, Fedyaeva AV, Varakina NN, Rusaleva TM, Borovskii GB, Rikhvanov EG. 2017. The role of flavin-containing enzymes in mitochondrial membrane hyperpolarization and ROS production in respiring *Saccharomyces cerevisiae* cells under heat-shock conditions. Sci Rep 7:14.

21. Suski J, Lebiedzinska M, Bonora M, Pinton P, Duszynski J, Wieckowski MR. 2018. Relation between mitochondrial membrane potential and ROS formation, p 357–381. In Palmeira CM, Moreno AJ (ed), Mitochondrial Bioenergetics: Methods and Protocols, 2nd Edition, vol 1782. Humana Press Inc, Totowa.

22. Malina C, Larsson C, Nielsen J. 2018. Yeast mitochondria: an overview of mitochondrial biology and the potential of mitochondrial systems biology. FEMS Yeast Res 18: foy040.

23. Melber A, Vashisht A, Weiler BD, Lill R, Wohlschlegel JA, Winge DR. 2016. Role of Nfu1 and Bol3 in iron-sulfur cluster transfer to mitochondrial clients. eLife 5:e15991.

24. Novodvorska M, Stratford M, Blythe MJ, Wilson R, Beniston RG, Archer DB. 2016. Metabolic activity in dormant conidia of *Aspergillus niger* and developmental changes during conidial outgrowth. Fungal Genet Biol 94:23–31.

25. Stratford M, Anslow PA. 1996. Comparison of the inhibitory action on *Saccharomyces cerevisiae* of weak-acid preservatives, uncouplers, and medium-chain fatty acids. FEMS Microbiol Lett 142:53–58.

26. Harada K, Higuchi R, Utsumi I. 1968. Studies on sorbic acid. 4. Inhibition of respiration in yeast. Agric Biol Chem 32:940–946.

27. Kusumegi K, Yoshida H, Tomiyama S. 1998. Inhibitory effects of acetic acid on respiration and growth of *Zygosaccharomyces rouxii*. J Ferment Bioeng 85:213–217.

28. York GK, Vaughn RH. 1964. Mechanisms in inhibition of microorganisms by sorbic acid. Journal of Bacteriology 88:411–417.

29. Islahudin F, Khozoie C, Bates S, Ting KN, Pleass RJ, Avery SV. 2013. Cell wall perturbation sensitizes fungi to the antimalarial drug chloroquine. Antimicrob Agents Chemother 57:3889–3896.

30. Winzeler EA, Shoemaker DD, Astromoff A, Liang H, Anderson K, Andre B, Bangham R, Benito R, Boeke JD, Bussey H, Chu AM, Connelly C, Davis K, Dietrich F, Dow SW, El Bakkoury M, Foury F, Friend SH, Gentalen E, Giaever G, Hegemann JH, Jones T, Laub M, Liao H, Liebundguth N, Lockhart DJ, Lucau-Danila A, Lussier M, M’Rabet N, Menard P, Mittmann M, Pai C, Rebischung C, Revuelta JL, Riles L, Roberts CJ, Ross-MacDonald P, Scherens B, Snyder M, Sookhai-Mahadeo S, Storms RK, Veronneau S, Voet M, Volckaert G, Ward TR, Wysocki R, Yen GS, Yu KX, Zimmermann K, Philippsen P, et al. 1999. Functional characterization of the *S. cerevisiae* genome by gene deletion and parallel analysis. Science 285:901–906.

31. Griffiths LM, Doudican NA, Shadel GS, Doetsch PW. 2009. Mitochondrial DNA oxidative damage and mutagenesis in *Saccharomyces cerevisiae*, p 267–286. In Stuart JA (ed), Mitochondrial DNA: Methods and Protocols, vol 554. Humana Press Inc, Totowa.

32. Imlay JA, Sethu R, Rohaun SK. 2019. Evolutionary adaptations that enable enzymes to tolerate oxidative stress. Free Rad Biol Med 140:4–13.

33. Vallieres C, Holland SL, Avery SV. 2017. Mitochondrial ferredoxin determines vulnerability of cells to copper excess. Cell Chem Biol 24:1228–1237.

34. den Besten HMW, Wells-Bennik MHJ, Zwietering MH. 2018. Natural diversity in heat resistance of bacteria and bacterial spores: Impact on food safety and quality, p 383–410. In Doyle MP, Klaenhammer TR (ed), Annual Review of Food Science and Technology, vol 9. Annual Reviews, Palo Alto.

35. Geoghegan IA, Stratford M, Bromley M, Archer DB, Avery SV. 2020. Weak Acid Resistance A (WarA), a novel transcription factor required for regulation of weak-acid resistance and spore-spore heterogeneity in *Aspergillus niger*. mSphere 5:e00685–19.

36. Stratford M, Steels H, Nebe-von-Caron G, Avery SV, Novodvorska M, Archer DB. 2014. Population heterogeneity and dynamics in starter culture and lag phase adaptation of the spoilage yeast *Zygosaccharomyces bailii* to weak acid preservatives. Int J Food Microbiol 181:40–47.

37. Stratford M, Steels H, Novodvorska M, Archer DB, Avery SV. 2019. Extreme osmotolerance and halotolerance in food-relevant yeasts and the role of glycerol-dependent cell individuality. Front Microbiol 9:14.

38. van Esch F. 1987. Yeasts in soft drinks and fruit juice concentrates. De Ware(n) Chemicus 17:20–31.

39. Chang AY, Marshall WF. 2017. Organelles - understanding noise and heterogeneity in cell biology at an intermediate scale. J Cell Sci 130:819–826.

40. Radzinski M, Fassler R, Yogev O, Breuer W, Shai N, Gutin J, Ilyas S, Geffen Y, Tsytkin-Kirschenzweig S, Nahmias Y, Ravid T, Friedman N, Shuldiner M, Reichmann D. 2018. Temporal profiling of redox-dependent heterogeneity in single cells. eLife 7:33.

41. Rodriguez A, Magan N, Medina A. 2016. Evaluation of the risk of fungal spoilage when substituting sucrose with commercial purified Stevia glycosides in sweetened bakery products. Int J Food Microbiol 231:42–47.

42. Kurtzman CP. 2003. Phylogenetic circumscription of *Saccharomyces, Kluyveromyces* and other members of the Saccharomycetaceae, and the proposal of the new genera *Lachancea, Nakaseomyces, Naumovia, Vanderwaltozyma* and *Zygotorulaspora*. FEMS Yeast Res 4:233–245.

43. Giorgini F, Guidetti P, Nguyen QV, Bennett SC, Muchowski PJ. 2005. A genomic screen in yeast implicates kynurenine 3-monooxygenase as a therapeutic target for Huntington disease. Nature Genet 37:526–531.

