## Supplementary material for "The preservative sorbic acid targets respiration, explaining the resistance of fermentative spoilage-yeast species": Suppl Tables S1, S3, S4, Fig S1

Stratford et al. - **SUPPLEMENTAL MATERIAL**

(TABLE S2 is provided in a separate file)

**TABLE S1** Principal yeast species used in this research. Original and current species names are provided (1) together with the strain sources.

| Strain | Previous Species | Current Species | Origin |
|---|---|---|---|
| NY40 | *Aureobasidium pullulans* | *Aureobasidium pullulans* | Factory airplate UK |
| 69 | *Candida parapsilosis* | *Candida parapsilosis* | Spoilage, fruit juice |
| 519 | *Candida pseudointermedia* | *Candida pseudointermedia* | Factory floor, Brazil |
| NCYC 3297 | *Candida pseudolambica* | *Candida pseudolambica* | Factory drain, Brazil |
| 546 | *Cryptococcus laurentii* | *Cryptococcus laurentii* | Factory drain, Brazil |
| 628 | *Cryptococcus magnus* | *Cryptococcus magnus* | Scrapper Factory, Russia |
| 522 | *Issatchenkia orientalis* | *Pichia kudriavzevii* | Factory drain, Brazil |
| NCYC 3371 | *Pichia anomala* | *Wicherhamomyces anomalus* | Factory, Israel |
| 92 | *Rhodotorula glutinis* | *Rhodotorula glutinis* | Factory, Israel |
| 95 | *Rhodotorula mucilaginosa* | *Rhodotorula mucilaginosa* | Factory wall, UK |
| NCYC 3368 | *Saccharomyces cerevisiae* | *Saccharomyces cerevisiae* | Wine yeast |
| BY4741 | *Saccharomyces cerevisiae* | *Saccharomyces cerevisiae* | Laboratory strain |
| BY4741 Δ*pad1* | *Saccharomyces cerevisiae* | *Saccharomyces cerevisiae* | Laboratory strain |
| BY4743 | *Saccharomyces cerevisiae* | *Saccharomyces cerevisiae* | Laboratory strain |
| BY4743 Δ*bol3* | *Saccharomyces cerevisiae* | *Saccharomyces cerevisiae* | Laboratory strain |
| BY4743 Δ*grx5* | *Saccharomyces cerevisiae* | *Saccharomyces cerevisiae* | Laboratory strain |
| BY4743 Δ*iba57* | *Saccharomyces cerevisiae* | *Saccharomyces cerevisiae* | Laboratory strain |
| BY4743 Δ*isu1* | *Saccharomyces cerevisiae* | *Saccharomyces cerevisiae* | Laboratory strain |
| BY4743 Δ*nfu1* | *Saccharomyces cerevisiae* | *Saccharomyces cerevisiae* | Laboratory strain |
| BY4743 Δ*ssq1* | *Saccharomyces cerevisiae* | *Saccharomyces cerevisiae* | Laboratory strain |
| 55 | *Saccharomyces exiguus* | *Kazachstania exigua* | Spoilage, mayonnaise salad |
| 529 | *Torulaspora delbruckii* | *Torulaspora delbruckii* | Factory filler, Brazil |
| 405 | *Trichosporon ovoides* | *Trichosporon ovoides* | Factory pallet, Turkey |
| NCYC 1766 | *Zygosaccharomyces bailii* | *Zygosaccharomyces bailii* | Spoilage, fruit juice |
| NCYC 1555 | *Zygosaccharomyces bisporus* | *Zygosaccharomyces bisporus* | Spoilage, salad cream |
| NCYC 2789 | *Zygosaccharomyces lentus* | *Zygosaccharomyces lentus* | Spoilage, orange juice |

**TABLE S3** Fermentation by spoilage and non-spoilage yeast species

| Group[a] | Strain | Yeast Species | Capacity to Ferment[b] | Glucose fermentation (mls/ml) | |
|---|---|---|---|---|---|
| | | | | 20 g/l[c] | 180 g/l |
| 3 | 628 | *Cryptococcus magnus* | No | 0[d] | 0 |
| 3 | 95 | *Rhodotorula mucilaginosa* | No | 0 | 0 |
| 3 | 92 | *Rhodotorula glutinis* | No | 0 | 0 |
| 3 | 546 | *Cryptococcus laurentii* | No | 0 | 0 |
| 2 | NCYC 3371 | *Wicherhamomyces anomalus* | Yes | 2.7 | 18.4 |
| 2 | 519 | *Candida pseudointermedia* | Yes | 2.8 | 11.6 |
| 2 | 69 | *Candida parapsilosis* | Yes | 2.8 | 9.6 |
| 2 | 529 | *Torulaspora delbruckii* | Yes | 3 | 34 |
| 2 | BY4741 | *Saccharomyces cerevisiae* | Yes | 3.2 | 34.5 |
| 2 | BY4741 Δ*pad1* | *Saccharomyces cerevisiae* | Yes | 3.3 | 36 |
| 2 | BY4741 petite | *Saccharomyces cerevisiae* | Yes | 3.1 | 35 |
| 1 | NCYC 3297 | *Candida pseudolambica* | Yes | 2.7 | 6.6 |
| 1 | 55 | *Kazachstania exigua* | Yes | 2.9 | 33.6 |
| 1 | 522 | *Pichia kudriavzevii* | Yes | 2.8 | 29 |
| 1 | NCYC 1555 | *Zygosaccharomyces bisporus* | Yes | 3.1 | 29.2 |
| 1 | NCYC 1766 | *Zygosaccharomyces bailii* | Yes | 3 | 36 |
| 1 | NCYC 2789 | *Zygosaccharomyces lentus* | Yes | 2.8 | 27.8 |

[a]Davenport grouping according to spoilage incidence (6). [b]According to (1). [c]Glucose concentration supplied. [d]Fermentation was determined according to the gas pressure after 28 days at 24°C in replicate, static bottles in YEP supplemented with the indicated glucose concentration.

**TABLE S4** Resistance of *S. cerevisiae* to weak acids with different carbon chain lengths.

| | Number of carbons | cLogP | Weak acid MIC[a] (mM) | | MIC ratio Glyc/Gluc |
|---|---|---|---|---|---|
| | | | Glucose | Glycerol | |
| Acetic acid | 2 | -0.194 | 130 | 117 | 90 |
| Propionic acid | 3 | 0.335 | 60 | 50 | 83 |
| Butyric acid | 4 | 0.864 | 26 | 21 | 81 |
| Valeric acid | 5 | 1.393 | 9.25 | 7 | 76 |
| Hexanoic acid | 6 | 1.922 | 3.1 | 2.1 | 68 |
| Heptanoic acid | 7 | 2.451 | 1.65 | 0.9 | 55 |
| Octanoic acid | 8 | 2.98 | 0.99 | 0.55 | 53 |

[a]Weak acid MIC, mean from replicate determinations after 14 d shaking at 120 rev. $min^{-1}$, 24°C in flasks containing YEP, pH 4.0 supplemented with weak acid and 30 g/l of either glucose or glycerol.

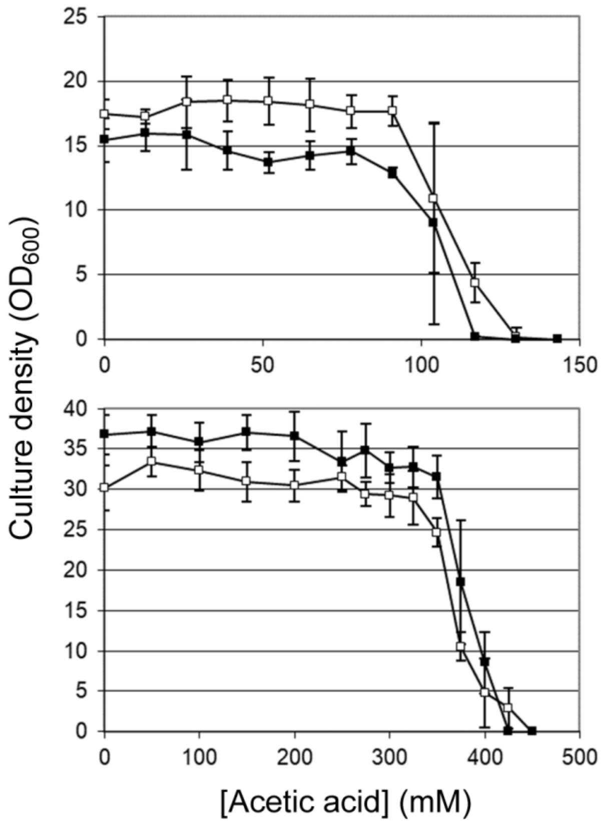


**FIG S1** Growth on glucose or glycerol in the presence of acetic acid. *S. cerevisiae* (top panel) or *Z. bailii* (bottom panel) were cultured in either 30 g/l glucose (open squares) or 30 g/l glycerol (closed squares), in YEP pH 4.0 supplemented with the indicated concentrations of acetic acid. $OD_{600}$ in flasks was determined after shaking at 120 rev. $min^{-1}$, 24°C for 14 days. Points are means from three replicate determinations ± S.D.
