## Supplementary material for "The preservative sorbic acid targets respiration, explaining the resistance of fermentative spoilage-yeast species": Suppl Table S2

**TABLE S2** Full list of yeast species, encompassing 687 strains, that were tested for fermentation and sorbic acid MIC (Figure 3). Strain numbers refer to the Mologic yeast collection.

| Yeast Species | Strain | Origin |
|---|---|---|
| *Barnettozyma californica* | 376 | Turkey |
| *Barnettozyma californica* | 538 | Brazil |
| *Barnettozyma sp.nov.* | 486 | Brazil |
| *Barnettozyma sp.nov.* | 497 | Brazil |
| *Barnettozyma subsufficiens* | 539 | Brazil |
| *Brettanomyces naardenensis* | 27 | UK |
| *Brettanomyces naardenensis* | 129 | UK |
| *Brettanomyces naardenensis* | 131 | UK |
| *Candida aaseri* | 440 | Turkey |
| *Candida aaseri* | 537 | Brazil |
| *Candida albicans* | 212 | Netherlands |
| *Candida albicans* | 309 | Italy |
| *Candida apicola* | 530 | Brazil |
| *Candida boidinii* | 166 | Netherlands |
| *Candida boidinii* | 204 | Netherlands |
| *Candida boidinii* | 295 | UK |
| *Candida boidinii* | 307 | Belgium |
| *Candida boidinii* | 368 | Turkey |
| *Candida boidinii* | 411 | Turkey |
| *Candida boidinii* | 420 | Turkey |
| *Candida boidinii* | 430 | Turkey |
| *Candida boidinii* | 464 | Russia |
| *Candida boidinii* | 487 | Brazil |
| *Candida boidinii* | 527 | Brazil |
| *Candida boidinii* | 557 | Brazil |
| *Candida boidinii* | 603 | Russia |
| *Candida boidinii* | 637 | UK |
| *Candida carpophila* | 350 | Thailand |
| *Candida davenportii* | 220 | Netherlands |
| *Candida diddensiae* | 355 | Thailand |
| *Candida diddensiae* | 553 | Brazil |
| *Candida diddensiae* | 611 | Russia |
| *Candida etchellsii* | 240 | Netherlands |
| *Candida inconspicua* sister sp. | 457 | Russia |
| *Candida intermedia* | 382 | Turkey |
| *Candida intermedia* | NCYC 2504 | |
| *Candida intermedia* | NCYC 2531 | |
| *Candida magnoliae* | M1 | UK |
| *Candida melibiosica* | 328 | Thailand |
| *Candida melibiosica* | 345 | Thailand |
| *Candida melibiosica* | 346 | Thailand |
| *Candida natalensis* | 518 | Brazil |

| Yeast Species | Strain | Origin |
|---|---|---|
| *Candida neerlandica* sister sp. | 493 | Brazil |
| *Candida norwegica* | 215 | Netherlands |
| *Candida oleophila* | 147 | Netherlands |
| *Candida oleophila* | 206 | Netherlands |
| *Candida oleophila* | 467 | Russia |
| *Candida oleophila* | 598 | Russia |
| *Candida orthopsilosis* | 238 | Thailand |
| *Candida orthopsilosis* | 358 | Thailand |
| *Candida orthopsilosis* | 517 | Brazil |
| *Candida parapsilosis* | 67 | UK |
| *Candida parapsilosis* | 68 | UK |
| *Candida parapsilosis* | 69 | UK |
| *Candida parapsilosis* | 138 | France |
| *Candida parapsilosis* | 157 | Netherlands |
| *Candida parapsilosis* | 163 | UK |
| *Candida parapsilosis* | 164 | UK |
| *Candida parapsilosis* | 203 | Ghana |
| *Candida parapsilosis* | 283 | UK |
| *Candida parapsilosis* | 384 | Turkey |
| *Candida parapsilosis* | 408 | Turkey |
| *Candida parapsilosis* | 432 | Turkey |
| *Candida parapsilosis* | 451 | Russia |
| *Candida parapsilosis* | 484 | Brazil |
| *Candida parapsilosis* | 583 | S Africa |
| *Candida parapsilosis* | 589 | Belgium |
| *Candida parapsilosis* | 625 | Russia |
| *Candida parapsilosis* | 630 | Germany |
| *Candida parapsilosis* | 644 | UK |
| *Candida parapsilosis* | 662 | UK |
| *Candida parapsilosis* | 672 | UK |
| *Candida parapsilosis* | 679 | UK |
| *Candida parapsilosis* | 682 | UK |
| *Candida pararugosa* | 122 | UK |
| *Candida pseudoglaebosa* | 223 | Netherlands |
| *Candida pseudointermedia* | 79 | France |
| *Candida pseudointermedia* | 153 | Netherlands |
| *Candida pseudointermedia* | 211 | Netherlands |
| *Candida pseudointermedia* | 227 | Netherlands |
| *Candida pseudointermedia* | 235 | Netherlands |
| *Candida pseudointermedia* | 246 | Netherlands |
| *Candida pseudointermedia* | 281 | France |
| *Candida pseudointermedia* | 327 | Thailand |
| *Candida pseudointermedia* | 332 | Thailand |
| *Candida pseudointermedia* | 340 | Thailand |
| *Candida pseudointermedia* | 351 | Thailand |
| *Candida pseudointermedia* | 372 | Turkey |
| *Candida pseudointermedia* | 407 | Turkey |
| *Candida pseudointermedia* | 416 | Turkey |
| *Candida pseudointermedia* | 419 | Turkey |

| Yeast Species | Strain | Origin |
|---|---|---|
| *Candida pseudointermedia* | 425 | Turkey |
| *Candida pseudointermedia* | 460 | Russia |
| *Candida pseudointermedia* | 495 | Brazil |
| *Candida pseudointermedia* | 496 | Brazil |
| *Candida pseudointermedia* | 501 | Brazil |
| *Candida pseudointermedia* | 519 | Brazil |
| *Candida pseudointermedia* | 528 | Brazil |
| *Candida pseudointermedia* | 552 | Brazil |
| *Candida pseudointermedia* | 556 | Brazil |
| *Candida pseudointermedia* | 590 | France |
| *Candida pseudointermedia* | 591 | France |
| *Candida pseudointermedia* | 606 | Russia |
| *Candida pseudointermedia* | 629 | Netherlands |
| *Candida pseudointermedia* | 639 | UK |
| *Candida pseudointermedia* | 640 | UK |
| *Candida pseudointermedia* | 680 | UK |
| *Candida pseudointermedia* | NCYC 2610 | |
| *Candida pseudointermedia* | NCYC 3278 | |
| *Candida pseudolambica* | 400 | Turkey |
| *Candida pseudolambica* | 435 | Turkey |
| *Candida pseudolambica* | 449 | Russia |
| *Candida pseudolambica* | 485 | Brazil |
| *Candida pseudolambica* | 525 | Brazil |
| *Candida pseudolambica* | 526 | Brazil |
| *Candida pseudolambica* | 599 | Russia |
| *Candida pseudolambica* | 605 | Russia |
| *Candida pseudolambica* | 618 | Russia |
| *Candida pseudolambica* | 627 | Russia |
| *Candida rugosa* | 353 | Thailand |
| *Candida sake* | 322 | UK |
| *Candida sake* | 323 | UK |
| *Candida silvae* | 626 | Russia |
| *Candida sojae* | 121 | UK |
| *Candida sojae* | 126 | France |
| *Candida sojae* | 165 | Poland |
| *Candida sojae* | 245 | Netherlands |
| *Candida sojae* | 312 | Brazil |
| *Candida sojae* | 375 | Turkey |
| *Candida sojae* | 414 | Turkey |
| *Candida sojae* | 434 | Turkey |
| *Candida sojae* | 439 | Turkey |
| *Candida sojae* | 446 | France |
| *Candida sojae* | 483 | Brazil |
| *Candida sojae* | 601 | Russia |
| *Candida sojae* | 608 | Russia |
| *Candida sorbophila* | 168 | Thailand |
| *Candida sorbophila* | NCYC 173 | |
| *Candida sorbosivorans* | 479 | Brazil |
| *Candida sp.* | 383 | Turkey |

| **Yeast Species** | **Strain** | **Origin** |
|---|---|---|
| *Candida sp.nov.* | 397 | China |
| *Candida sp.nov.* | 452 | Russia |
| *Candida sp.nov.* | 524 | Brazil |
| *Candida sp.nov.* | 577 | Germany |
| *Candida sp.nov.* | 674 | UK |
| *Candida sp.nov.* | 675 | UK |
| *Candida sp.nov.* | 681 | UK |
| *Candida sp.nov.* | 683 | UK |
| *Candida tropicalis* | 342 | Thailand |
| *Candida tropicalis* | 344 | Thailand |
| *Candida tropicalis* | 352 | Thailand |
| *Candida tropicalis* | 369 | Turkey |
| *Candida tropicalis* | 389 | Turkey |
| *Candida tropicalis* | 417 | Turkey |
| *Candida tropicalis* | 428 | Turkey |
| *Candida tropicalis* | 520 | Brazil |
| *Candida tropicalis* sister sp. | 377 | Turkey |
| *Candida tropicalis* sister sp. | 492 | Brazil |
| *Candida vartiovaarae* | 622 | Russia |
| *Candida versitalis* | 396 | China |
| *Candida wyomingensis* | 617 | Russia |
| *Candida zeylandoides* | 26 | UK |
| *Clavispora lusitania* | 53 | Israel |
| *Clavispora lusitania* | 74 | France |
| *Clavispora lusitania* | 110 | Argentina |
| *Clavispora lusitania* | 225 | Netherlands |
| *Clavispora lusitania* | 399 | Germany |
| *Clavispora lusitania* | 436 | Turkey |
| *Clavispora lusitania* | 443 | Turkey |
| *Clavispora lusitania* | 444 | Turkey |
| *Clavispora lusitania* | 445 | Turkey |
| *Clavispora lusitania* | 461 | Russia |
| *Clavispora lusitania* | 462 | Russia |
| *Clavispora lusitania* | 481 | Brazil |
| *Clavispora lusitania* | 531 | Brazil |
| *Clavispora lusitania* | 559 | Brazil |
| *Clavispora lusitania* | 643 | UK |
| *Clavispora lusitania* | 655 | UK |
| *Clavispora lusitania* | 671 | UK |
| *Clavispora lusitania* | 53a | Israel |
| *Clavispora lusitania* | NCYC 3268 | |
| *Cryptococcus albidosimilis* | 361 | Turkey |
| *Cryptococcus albidus* | 144 | Netherlands |
| *Cryptococcus albidus* | 216 | Netherlands |
| *Cryptococcus albidus* | 660 | UK |
| *Cryptococcus albidus* | 673 | UK |
| *Cryptococcus albidus* | NCYC 445 | |
| *Cryptococcus cylindricus* | 82 | UK |
| *Cryptococcus diffluens* | 600 | Russia |

| **Yeast Species** | **Strain** | **Origin** |
|---|---|---|
| *Cryptococcus flavescens* | 214 | Netherlands |
| *Cryptococcus humicola* | 490 | Brazil |
| *Cryptococcus laurentii* | 60 | UK |
| *Cryptococcus laurentii* | 77 | France |
| *Cryptococcus laurentii* | 78 | France |
| *Cryptococcus laurentii* | 99 | UK |
| *Cryptococcus laurentii* | 341 | Thailand |
| *Cryptococcus laurentii* | 347 | Thailand |
| *Cryptococcus laurentii* | 360 | Turkey |
| *Cryptococcus laurentii* | 433 | Turkey |
| *Cryptococcus laurentii* | 480 | Brazil |
| *Cryptococcus laurentii* | 546 | Brazil |
| *Cryptococcus laurentii* | 607 | Russia |
| *Cryptococcus laurentii* | 609 | Russia |
| *Cryptococcus laurentii* sister sp | 404 | Turkey |
| *Cryptococcus laurentii* sister sp | 580 | S Africa |
| *Cryptococcus laurentii* sister sp | 664 | UK |
| *Cryptococcus liquefaciens* | 349 | Thailand |
| *Cryptococcus liquefaciens* | 357 | Thailand |
| *Cryptococcus liquefaciens* | 402 | Turkey |
| *Cryptococcus liquefaciens* | 418 | Turkey |
| *Cryptococcus liquefaciens* | 424 | Turkey |
| *Cryptococcus magnus* | 422 | Turkey |
| *Cryptococcus magnus* | 465 | Russia |
| *Cryptococcus magnus* | 628 | Russia |
| *Cryptococcus magnus* | 645 | UK |
| *Cryptococcus magnus* | 666 | UK |
| *Cryptococcus nyarrowii* sister sp. | 85 | UK |
| *Cryptococcus ramirezgomezianus* | 145 | Netherlands |
| *Cryptococcus saitou* | 234 | Netherlands |
| *Cryptococcus saitou* | 448 | Russia |
| *Cryptococcus saitou* | 584 | S Africa |
| *Cryptococcus saitou* | 663 | UK |
| *Cryptococcus sp.nov.* | 84 | UK |
| *Cryptococcus sp.nov.* | 547 | Brazil |
| *Cryptococcus sp.nov.* | 548 | Brazil |
| *Cryptococcus uzbekistanensis* | 586 | S Africa |
| *Cryptococcus victoriae* sister sp. | 665 | UK |
| *Debaryomyces hansenii* | 146 | Netherlands |
| *Debaryomyces hansenii* | 403 | Turkey |
| *Debaryomyces hansenii* | 437 | Turkey |
| *Debaryomyces hansenii* | 450 | Russia |
| *Debaryomyces hansenii* | 657 | UK |
| *Debaryomyces hansenii* | NCYC 9 | |
| *Debaryomyces hansenii* var *fabryii* | 100 | UK |
| *Debaryomyces hansenii* var *fabryii* | 558 | Brazil |
| *Dekkera anomala* | 247 | Belgium |
| *Dekkera anomala* | 502 | UK |
| *Dekkera anomala* | 506 | UK |

| **Yeast Species** | **Strain** | **Origin** |
|---|---|---|
| *Dekkera anomala* | 247 | Belgium |
| *Dekkera anomala* | 506 | UK |
| *Dekkera bruxellensis* | 148 | Netherlands |
| *Dekkera bruxellensis* | 306 | Belgium |
| *Dekkera bruxellensis* | 311 | UK |
| *Dekkera bruxellensis* | 319 | Belgium |
| *Dekkera bruxellensis* | 325 | Belgium |
| *Dekkera bruxellensis* | 326 | Belgium |
| *Dekkera bruxellensis* | 507 | UK |
| *Dekkera bruxellensis* | NCYC 823 | |
| *Filobasidiella neoformans* | 545 | Brazil |
| *Filobasidium uniguttulatum* | 230 | Netherlands |
| *Hanseniaspora guilliermondii* | NRRL 1625 | |
| *Hanseniaspora meyeri* | 81 | UK |
| *Hanseniaspora meyeri* | 127 | France |
| *Hanseniaspora meyeri* | 155 | Netherlands |
| *Hanseniaspora occidentalis* | NRRL 7946 | |
| *Hanseniaspora osmophila* | NRRL 1613 | |
| *Hanseniaspora uvarum* | 226 | Netherlands |
| *Hanseniaspora uvarum* | 321 | UK |
| *Hanseniaspora uvarum* | 646 | UK |
| *Hanseniaspora uvarum* | NRRL 1614 | |
| *Hanseniaspora uvarum* | NRRL 1626 | |
| *Hanseniaspora vineae* | NRRL17529 | |
| *Kazachstania barnettii* | 57 | UK |
| *Kazachstania exigua* | 23 | UK |
| *Kazachstania exigua* | 55 | UK |
| *Kazachstania exigua* | 152 | Netherlands |
| *Kazachstania kunashirensis* | NCYC 2702 | |
| *Kazachstania martiniae* | NCYC 2703 | |
| *Kazachstania servazzii* | 58 | UK |
| *Kazachstania servazzii* | NCYC 2577 | |
| *Kazachstania unispora* | NCYC 971 | |
| *Kloeckera linderi* | NRRL17531 | |
| *Kluyveromyces marxianus* | 562 | UK |
| *Kodamaea ohmeri* | 348 | Thailand |
| *Kodamaea ohmeri* | 535 | Brazil |
| *Komagataella pastoris* | 219 | Netherlands |
| *Kregervanrija fluxuum* | 504 | UK |
| *Lachancea cidri* | NCYC 2875 | |
| *Lachancea fermentati* | NCYC 2508 | |
| *Lindnera fabianii* | 534 | Brazil |
| *Lindnera jadinii* | 365 | Turkey |
| *Lindnera jadinii* | 477 | Brazil |
| *Lindnera jadinii* | 604 | Russia |
| *Lindnera jadinii* sister sp. | 209 | Netherlands |
| *Lodderomyces elongisporus* | 373 | Turkey |
| *Metschnikowia sp.nov.* | 229 | Netherlands |
| *Metschnikowia sp.nov.* | 536 | Brazil |

| Yeast Species | Strain | Origin |
|---|---|---|
| *Meyerozyma guilliermondii* | 224 | Netherlands |
| *Meyerozyma guilliermondii* | 343 | Thailand |
| *Meyerozyma guilliermondii* | 370 | Turkey |
| *Meyerozyma guilliermondii* | 374 | Turkey |
| *Meyerozyma guilliermondii* | 395 | Germany |
| *Meyerozyma guilliermondii* | 406 | Turkey |
| *Meyerozyma guilliermondii* | 427 | Turkey |
| *Meyerozyma guilliermondii* | 453 | Russia |
| *Meyerozyma guilliermondii* | 532 | Brazil |
| *Meyerozyma guilliermondii* | 620 | Russia |
| *Meyerozyma guilliermondii* | 659 | UK |
| *Meyerozyma guilliermondii* | M4 | UK |
| *Millerozyma farinosa* | 356 | Thailand |
| *Nakaseomyces glabrata* | 670 | UK |
| *Nakazawaea holstii* | 610 | Russia |
| *Pichia fermentans* | 293 | UK |
| *Pichia fermentans* | 324 | UK |
| *Pichia kudriavzevii* | 294 | UK |
| *Pichia kudriavzevii* | 172 | Netherlands |
| *Pichia kudriavzevii* | 217 | Netherlands |
| *Pichia kudriavzevii* | 354 | Thailand |
| *Pichia kudriavzevii* | 380 | Turkey |
| *Pichia kudriavzevii* | 522 | Brazil |
| *Pichia kudriavzevii* | 641 | UK |
| *Pichia kudriavzevii* | 654 | UK |
| *Pichia manshurica* | 86 | Netherlands |
| *Pichia manshurica* | 120 | UK |
| *Pichia manshurica* | 123 | UK |
| *Pichia manshurica* | 124 | UK |
| *Pichia manshurica* | 169 | Netherlands |
| *Pichia manshurica* | 170 | Netherlands |
| *Pichia manshurica* | 171 | Netherlands |
| *Pichia manshurica* | 236 | Thailand |
| *Pichia manshurica* | 386 | Turkey |
| *Pichia manshurica* | 458 | Russia |
| *Pichia manshurica* | 478 | Brazil |
| *Pichia manshurica* | 521 | Brazil |
| *Pichia membranifaciens* | 173 | UK |
| *Pichia membranifaciens* | 210 | Netherlands |
| *Pichia occidentalis* | 237 | Thailand |
| *Pichia occidentalis* | 454 | Russia |
| *Pichia occidentalis* | 473 | Brazil |
| *Pichia occidentalis* | 523 | Brazil |
| *Pichia scutulata* sister sp. | 320 | UK |
| *Pichia sp.nov.* | 202 | UK |
| *Pseudozyma aphidis* | 554 | Brazil |
| *Pseudozyma sp.nov.* | 336 | Thailand |
| *Pseudozyma sp.nov.* | 658 | UK |
| *Rhodosporidium fluviale* | 379 | Turkey |

| **Yeast Species** | **Strain** | **Origin** |
|---|---|---|
| *Rhodosporidium fluviale* | 426 | Turkey |
| *Rhodosporidium fluviale* | 491 | Brazil |
| *Rhodosporidium fluviale* | 549 | Brazil |
| *Rhodotorula colostri* | 623 | Russia |
| *Rhodotorula dairenensis* | 616 | Russia |
| *Rhodotorula glutinis* | 92 | Israel |
| *Rhodotorula glutinis* | 96 | France |
| *Rhodotorula glutinis* | 330 | Thailand |
| *Rhodotorula glutinis* | 412 | Turkey |
| *Rhodotorula glutinis* sister sp. | NCYC 59 | |
| *Rhodotorula glutinis* sister sp. | 93 | France |
| *Rhodotorula graminis* | 167 | Thailand |
| *Rhodotorula graminis* | 587 | S Africa |
| *Rhodotorula graminis* | 615 | Russia |
| *Rhodotorula minuta* | 378 | Turkey |
| *Rhodotorula mucilaginosa* | 90 | UK |
| *Rhodotorula mucilaginosa* | 95 | UK |
| *Rhodotorula mucilaginosa* | 143 | Netherlands |
| *Rhodotorula mucilaginosa* | 218 | Netherlands |
| *Rhodotorula mucilaginosa* | 222 | Netherlands |
| *Rhodotorula mucilaginosa* | 329 | Thailand |
| *Rhodotorula mucilaginosa* | 363 | Turkey |
| *Rhodotorula mucilaginosa* | 401 | Turkey |
| *Rhodotorula mucilaginosa* | 441 | Turkey |
| *Rhodotorula mucilaginosa* | 468 | Russia |
| *Rhodotorula mucilaginosa* | 469 | Russia |
| *Rhodotorula mucilaginosa* | 489 | Brazil |
| *Rhodotorula mucilaginosa* | 550 | Brazil |
| *Rhodotorula mucilaginosa* | 581 | S Africa |
| *Rhodotorula mucilaginosa* | 668 | UK |
| *Rhodotorula mucilaginosa* | 676 | UK |
| *Rhodotorula mucilaginosa* | 677 | UK |
| *Rhodotorula mucilaginosa* | NCYC 195 | |
| *Rhodotorula nothofagi* | 154 | Netherlands |
| *Rhodotorula nothofagi* | 602 | Russia |
| *Rhodotorula slooffiae* | 208 | Netherlands |
| *Rhodotorula slooffiae* | 585 | S Africa |
| *Rhodotorula sp.nov.* | 101 | UK |
| *Saccharomyces bayanus* var *bayanus* | 24 | UK |
| *Saccharomyces bayanus* var *bayanus* | 59 | UK |
| *Saccharomyces bayanus* var *bayanus* | NCYC 2669 | |
| *Saccharomyces bayanus* var *uvarum* | 97 | UK |
| *Saccharomyces bayanus* var *uvarum* | 25 | UK |
| *Saccharomyces bayanus* var *uvarum* | 117 | UK |
| *Saccharomyces cariocanus* | NCYC 2890 | |
| *Saccharomyces cerevisiae* | 22 | UK |
| *Saccharomyces cerevisiae* | 47 | UK |
| *Saccharomyces cerevisiae* | 48 | UK |
| *Saccharomyces cerevisiae* | 56 | UK |

| **Yeast Species** | **Strain** | **Origin** |
|---|---|---|
| *Saccharomyces cerevisiae* | 62 | UK |
| *Saccharomyces cerevisiae* | 63 | UK |
| *Saccharomyces cerevisiae* | 64 | UK |
| *Saccharomyces cerevisiae* | 65 | UK |
| *Saccharomyces cerevisiae* | 125 | France |
| *Saccharomyces cerevisiae* | 174 | UK |
| *Saccharomyces cerevisiae* | 244 | Netherlands |
| *Saccharomyces cerevisiae* | 253 | Netherlands |
| *Saccharomyces cerevisiae* | 273 | UK |
| *Saccharomyces cerevisiae* | 282 | Netherlands |
| *Saccharomyces cerevisiae* | 291 | UK |
| *Saccharomyces cerevisiae* | 292 | UK |
| *Saccharomyces cerevisiae* | 308 | Belgium |
| *Saccharomyces cerevisiae* | 317 | UK |
| *Saccharomyces cerevisiae* | 359 | Turkey |
| *Saccharomyces cerevisiae* | 632 | UK |
| *Saccharomyces cerevisiae* | 633 | UK |
| *Saccharomyces cerevisiae* | 634 | UK |
| *Saccharomyces cerevisiae* | 635 | UK |
| *Saccharomyces cerevisiae* | 636 | UK |
| *Saccharomyces cerevisiae* | 656 | UK |
| *Saccharomyces cerevisiae* | 667 | UK |
| *Saccharomyces cerevisiae* | BY4741 | Euroscarf |
| *Saccharomyces cerevisiae* | BY4742 | Euroscarf |
| *Saccharomyces cerevisiae* | BY4743 | Euroscarf |
| *Saccharomyces cerevisiae* | NCYC 366 | |
| *Saccharomyces cerevisiae* | NCYC 87 | |
| *Saccharomyces cerevisiae* | X2180-1B | |
| *Saccharomyces kudriavzevii* | NCYC 2889 | |
| *Saccharomyces mikatae* | NCYC 2888 | |
| *Saccharomyces paradoxus* | NCYC 2600 | |
| *Saccharomyces paradoxus* | NCYC 2601 | |
| *Saccharomyces pastorianus* | NCYC 392 | |
| *Saccharomyces pastorianus* | 201 | Netherlands |
| *Saccharomycodes ludwigii* | NCYC 3532 | |
| *Saccharomycodes ludwigii* | NCYC 730 | |
| *Saccharomycodes ludwigii* | NCYC 731 | |
| *Saccharomycodes ludwigii* | NCYC 732 | |
| *Saccharomycodes ludwigii* | NCYC 734 | |
| *Saccharomycodes ludwigii* | NCYC 849 | |
| *Saturnispora sp.nov.* | 186 | UK |
| *Schizosaccharomyces pombe* | NCYC 1346 | |
| *Schizosaccharomyces pombe* | NCYC 2722 | |
| *Schwanniomyces etchelsii* | 213 | Netherlands |
| *Sporidiobolus johnsonii* | 498 | Brazil |
| *Sporidiobolus metaroseus* | 648 | UK |
| *Sporidiobolus metaroseus* | 649 | UK |
| *Sporidiobolus salmonicolor* | 221 | Netherlands |
| *Sporobolomyces sp.nov.* | 614 | Russia |

| **Yeast Species** | **Strain** | **Origin** |
|---|---|---|
| *Torulaspora delbrueckii* | 137 | UK |
| *Torulaspora delbrueckii* | 207 | Netherlands |
| *Torulaspora delbrueckii* | 366 | Turkey |
| *Torulaspora delbrueckii* | 410 | Turkey |
| *Torulaspora delbrueckii* | 456 | Russia |
| *Torulaspora delbrueckii* | 529 | Brazil |
| *Torulaspora delbrueckii* | 624 | Russia |
| *Torulaspora delbrueckii* | 631 | UK |
| *Torulaspora delbrueckii* | 638 | UK |
| *Torulaspora delbrueckii* | 653 | UK |
| *Torulaspora globosa* | NCYC 820 | |
| *Torulaspora microellipsoides* | NCYC 2568 | |
| *Torulaspora microellipsoides* | M11 | UK |
| *Torulaspora microellipsoides* | NCYC 411 | |
| *Torulaspora pretoriensis* | NCYC 524 | |
| *Trichosporon asahii* | 331 | Thailand |
| *Trichosporon asahii* | 429 | Turkey |
| *Trichosporon asahii* | 488 | Brazil |
| *Trichosporon asahii* | 540 | Brazil |
| *Trichosporon coremiiforme* | 459 | Russia |
| *Trichosporon coremiiforme* | 650 | UK |
| *Trichosporon coremiiforme* | 669 | UK |
| *Trichosporon coremiiforme* | 678 | UK |
| *Trichosporon domesticum* | 651 | UK |
| *Trichosporon gracile* | 647 | UK |
| *Trichosporon jirovecii* | 405 | Turkey |
| *Trichosporon jirovecii* | 542 | Brazil |
| *Trichosporon jirovecii* | 652 | UK |
| *Trichosporon mucoides* | 381 | Turkey |
| *Trichosporon mucoides* | 466 | Russia |
| *Trichosporon mycotoxinivorans* | 541 | Brazil |
| *Trichosporon ovoides* | 409 | Turkey |
| *Trichosporon sp.nov.* | 339 | Thailand |
| *Wickerhamomyces anomalus* | 54 | France |
| *Wickerhamomyces anomalus* | 70 | Israel |
| *Wickerhamomyces anomalus* | 71 | France |
| *Wickerhamomyces anomalus* | 73 | France |
| *Wickerhamomyces anomalus* | 88 | UK |
| *Wickerhamomyces anomalus* | 156 | Netherlands |
| *Wickerhamomyces anomalus* | 364 | Turkey |
| *Wickerhamomyces anomalus* | 415 | Turkey |
| *Wickerhamomyces anomalus* | 447 | Turkey |
| *Wickerhamomyces anomalus* | 455 | Russia |
| *Wickerhamomyces anomalus* | 482 | Brazil |
| *Wickerhamomyces anomalus* | 516 | Brazil |
| *Wickerhamomyces anomalus* | 582 | S Africa |
| *Wickerhamomyces anomalus* | 612 | Russia |
| *Wickerhamomyces anomalus* | 613 | Russia |
| *Wickerhamomyces anomalus* | NCYC 18 | |

| **Yeast Species** | **Strain** | **Origin** |
|---|---|---|
| *Wickerhamomyces anomalus* | NCYC 711 | |
| *Wickerhamomyces subpelliculosus* | IFFI 01014 | |
| *Yarrowia lipolytica* | 149 | Netherlands |
| *Yarrowia lipolytica* | 205 | Netherlands |
| *Yarrowia lipolytica* | 472 | Brazil |
| *Yarrowia lipolytica* | 474 | Brazil |
| *Yarrowia lipolytica* | 499 | Brazil |
| *Yarrowia lipolytica* | 500 | Brazil |
| *Yarrowia lipolytica* | 560 | Brazil |
| *Yarrowia lipolytica* | 619 | Russia |
| *Yarrowia lipolytica* | 642 | UK |
| *Yarrowia lipolytica* | 661 | UK |
| *Zygoascus hellenicus* | 533 | Brazil |
| *Zygosaccharomyces bailii* | 2 | UK |
| *Zygosaccharomyces bailii* | 4 | USA |
| *Zygosaccharomyces bailii* | 5 | USA |
| *Zygosaccharomyces bailii* | 6 | USA |
| *Zygosaccharomyces bailii* | 7 | USA |
| *Zygosaccharomyces bailii* | 8 | USA |
| *Zygosaccharomyces bailii* | 9 | USA |
| *Zygosaccharomyces bailii* | 10 | USA |
| *Zygosaccharomyces bailii* | 11 | USA |
| *Zygosaccharomyces bailii* | 12 | USA |
| *Zygosaccharomyces bailii* | 13 | USA |
| *Zygosaccharomyces bailii* | 15 | Netherlands |
| *Zygosaccharomyces bailii* | 16 | Netherlands |
| *Zygosaccharomyces bailii* | 17 | UK |
| *Zygosaccharomyces bailii* | 18 | UK |
| *Zygosaccharomyces bailii* | 19 | UK |
| *Zygosaccharomyces bailii* | 20 | UK |
| *Zygosaccharomyces bailii* | 21 | UK |
| *Zygosaccharomyces bailii* | 52 | Netherlands |
| *Zygosaccharomyces bailii* | 80 | Mexico |
| *Zygosaccharomyces bailii* | 105 | UK |
| *Zygosaccharomyces bailii* | 106 | UK |
| *Zygosaccharomyces bailii* | 107 | UK |
| *Zygosaccharomyces bailii* | 108 | UK |
| *Zygosaccharomyces bailii* | 112 | Belgium |
| *Zygosaccharomyces bailii* | 114 | Belgium |
| *Zygosaccharomyces bailii* | 119 | Netherlands |
| *Zygosaccharomyces bailii* | 194 | USA |
| *Zygosaccharomyces bailii* | 280 | S Africa |
| *Zygosaccharomyces bailii* | 362 | Turkey |
| *Zygosaccharomyces bailii* | 475 | Brazil |
| *Zygosaccharomyces bailii* | 503 | UK |
| *Zygosaccharomyces bailii* | 505 | UK |
| *Zygosaccharomyces bailii* | 592 | Phillipines |
| *Zygosaccharomyces bailii* | 593 | Phillipines |
| *Zygosaccharomyces bailii* | 594 | Sweden |

| **Yeast Species** | **Strain** | **Origin** |
|---|---|---|
| *Zygosaccharomyces bailii* | 595 | Spain |
| *Zygosaccharomyces bailii* | M10 | UK |
| *Zygosaccharomyces bailii* | M5 | UK |
| *Zygosaccharomyces bailii* | M6 | UK |
| *Zygosaccharomyces bailii* | M7 | UK |
| *Zygosaccharomyces bailii* | M8 | UK |
| *Zygosaccharomyces bailii* | NCYC 1416 | |
| *Zygosaccharomyces bailii* | NCYC 1766 | |
| *Zygosaccharomyces bisporus* | 28 | Israel |
| *Zygosaccharomyces bisporus* | 104 | UK |
| *Zygosaccharomyces bisporus* | 133 | UK |
| *Zygosaccharomyces bisporus* | 134 | UK |
| *Zygosaccharomyces bisporus* | 257 | Netherlands |
| *Zygosaccharomyces bisporus* | 367 | Turkey |
| *Zygosaccharomyces bisporus* | 390 | Turkey |
| *Zygosaccharomyces bisporus* | 391 | Turkey |
| *Zygosaccharomyces bisporus* | 494 | Brazil |
| *Zygosaccharomyces bisporus* | NCYC 1495 | |
| *Zygosaccharomyces bisporus* | NCYC 171 | |
| *Zygosaccharomyces bisporus* | NRRL 1228 | |
| *Zygosaccharomyces bisporus* | NRRL12627 | |
| *Zygosaccharomyces bisporus* | NRRL 7253 | |
| *Zygosaccharomyces bisporus* | NRRL 7684 | |
| *Zygosaccharomyces kombuchaensis* | 198 | USA |
| *Zygosaccharomyces kombuchaensis* | 199 | USA |
| *Zygosaccharomyces kombuchaensis* | 200 | USA |
| *Zygosaccharomyces kombuchaensis* | NCYC 2969 | |
| *Zygosaccharomyces lentus* | 36 | UK |
| *Zygosaccharomyces lentus* | 37 | UK |
| *Zygosaccharomyces lentus* | 38 | UK |
| *Zygosaccharomyces lentus* | 39 | France |
| *Zygosaccharomyces lentus* | 40 | UK |
| *Zygosaccharomyces lentus* | 103 | UK |
| *Zygosaccharomyces lentus* | 398 | UK |
| *Zygosaccharomyces lentus* | M9 | UK |
| *Zygosaccharomyces lentus* | TNO 0566 | |
| *Zygosaccharomyces lentus* | TNO 0567 | |
| *Zygosaccharomyces lentus* | TNO 0569 | |
| *Zygosaccharomyces lentus* | TNO 0572 | |
| *Zygosaccharomyces mellis* | 139 | UK |
| *Zygosaccharomyces mellis* | 141 | UK |
| *Zygosaccharomyces mellis* | 142 | UK |
| *Zygosaccharomyces mellis* | 192 | USA |
| *Zygosaccharomyces mellis* | 193 | USA |
| *Zygosaccharomyces mellis* | 195 | USA |
| *Zygosaccharomyces mellis* | 196 | USA |
| *Zygosaccharomyces mellis* | 197 | USA |
| *Zygosaccharomyces mellis* | NCYC 2403 | |
| *Zygosaccharomyces rouxii* | 33 | UK |

| **Yeast Species** | **Strain** | **Origin** |
|---|---|---|
| *Zygosaccharomyces rouxii* | 34 | UK |
| *Zygosaccharomyces rouxii* | 35 | UK |
| *Zygosaccharomyces rouxii* | 115 | UK |
| *Zygosaccharomyces rouxii* | 116 | UK |
| *Zygosaccharomyces rouxii* | 140 | Germany |
| *Zygosaccharomyces rouxii* | 231 | Denmark |
| *Zygosaccharomyces rouxii* | 232 | Denmark |
| *Zygosaccharomyces rouxii* | 233 | Denmark |
| *Zygosaccharomyces rouxii* | 239 | Netherlands |
| *Zygosaccharomyces rouxii* | 241 | Netherlands |
| *Zygosaccharomyces rouxii* | 254 | Netherlands |
| *Zygosaccharomyces rouxii* | 255 | Netherlands |
| *Zygosaccharomyces rouxii* | 265 | Netherlands |
| *Zygosaccharomyces rouxii* | 596 | Spain |
| *Zygosaccharomyces rouxii* | 597 | Spain |
| *Zygosaccharomyces rouxii* | ATCC66069 | |
| *Zygosaccharomyces rouxii* | CBS 4021 | |
| *Zygosaccharomyces rouxii* | CBS 4837 | |
| *Zygosaccharomyces rouxii* | CBS 681 | |
| *Zygosaccharomyces rouxii* | IFFI 01378 | |
| *Zygosaccharomyces rouxii* | IFFI 01417 | |
| *Zygosaccharomyces rouxii* | IFFI 01708 | |
| *Zygosaccharomyces rouxii* | M2 | UK |
| *Zygosaccharomyces rouxii* | M3 | UK |
| *Zygosaccharomyces rouxii* | NCYC 381 | |
| *Zygosaccharomyces rouxii* | NCYC 568 | |
| *Zygosaccharomyces rouxii* | NCYC 579 | |
| *Zygosaccharomyces rouxii* | NRRL 2547 | |
| *Zygosaccharomyces rouxii hybrid* | 252 | Netherlands |
| *Zygosaccharomyces rouxii hybrid* | 258 | Netherlands |
| *Zygosaccharomyces rouxii hybrid* | 259 | Netherlands |
| *Zygosaccharomyces rouxii hybrid* | 266 | Netherlands |
| *Zygosaccharomyces rouxii hybrid* | ATCC13356 | |
| *Zygosaccharomyces rouxii hybrid* | ATCC46261 | |
| *Zygosaccharomyces rouxii hybrid* | IFFI 01379 | |
| *Zygosaccharomyces rouxii hybrid* | IFFI 01711 | |
| *Zygosaccharomyces rouxii hybrid* | IFFI 01712 | |
| *Zygosaccharomyces rouxii hybrid* | NCYC 3363 | |
| *Zygosaccharomyces rouxii hybrid* | NRRL 2547 | |
| *Zygosaccharomyces sp.nov.* | IFFI 01710 | |
| *Zygosaccharomyces sp.nov.* | IFFI 01709 | |
| *Zygosaccharomyces sp.nov.* | NCYC 3265 | |
| *Zygotorulaspora florentina* | NCYC 2513 | |
| *Zygotorulaspora florentina* | 44 | UK |
| *Zygotorulaspora mrakii* | NCYC 2489 | |
